# Melanocyte differentiation and mechanosensation are differentially modulated by distinct extracellular matrix proteins

**DOI:** 10.1101/2024.10.04.616635

**Authors:** Carole Luthold, Marie Didion, Vanessa Samira Rácz, Emilio Benedum, Ann-Kathrin Burkhart, Nina Demmerle, Evelyn Wirth, Gubesh Gunaratnam, Sudharshini Thangamurugan, Volkhard Helms, Markus Bischoff, Annika Ridzal, Sandra Iden

**Author notes:** **CONTACT DETAILS** CL, SI: Cell & Developmental Biology, Saarland University, Faculty of Medicine, Kirrberger Strasse, Building 61.4, 66421 Homburg/Saar, Germany.

## Abstract

Dysfunctions in melanocytes can lead to pigmentation disorders, such as albinism, or contribute to the development of melanoma, the most aggressive form of skin cancer. Epidermal melanocytes typically interact with the collagen IV-rich basement membrane, but upon injury or in pathological conditions, they can encounter environments rich in collagen I or fibronectin. While alterations in extracellular matrix (ECM) composition and stiffness are known to impact cell behavior, the specific roles of each of these cues for melanocyte functions remain unclear. To explore the impact of these extrinsic cues, we here exposed murine melanocytes to different ECM proteins as well as varying substrate stiffnesses. This study identified MITF, a key regulator of melanocyte differentiation and function, as an ECM- and mechanosensitive transcription factor. We further revealed that distinct ECM proteins and substrate stiffness engage a FAK/MEK/ERK/MITF signaling axis to control melanocyte functions. Exposure of melanocytes to collagen I restricted FAK and ERK activation, promoting high nuclear MITF levels associated with melanocyte proliferation and differentiation. Conversely, fibronectin elicited elevated FAK and ERK activation, leading to low nuclear MITF, correlating with a dedifferentiated and motile phenotype. Consistent with these observations, RNA sequencing revealed that collagen I supports a differentiated gene expression program, whereas fibronectin induces a dedifferentiated transcriptomic signature. Importantly, inhibiting MEK or ERK activity in melanocytes cultured on fibronectin led to increased MITF nuclear localization and enhanced melanogenesis. Additionally, on fibronectin FAK inhibition reduced ERK activation and enhanced melanogenesis, supporting a role for FAK upstream of ERK. Finally, we uncovered that melanocyte mechanoresponses differ depending on the specific ECM environment. Together, these findings reveal a substantial effect of extrinsic cues on melanocyte function, with a context-dependent MITF regulation downstream of ERK, offering new perspectives for our understanding of melanocyte-related pathologies.

## INTRODUCTION

Melanocytes (MCs) are pigment-producing cells found in the basal layer of the skin epidermis, and in various organs such as the brain, heart, and eyes (Centeno et al., 2023). The pronounced dendritic morphology of cutaneous MCs enables an efficient melanosome transfer to surrounding keratinocytes, providing photoprotection (Benito-Martínez et al., 2021; Cui & Man, 2023; Domingues et al., 2020; Hirobe, 2014; Prospéri et al., 2024). Dysfunctions of MCs can cause pigmentation disorders such as albinism and vitiligo (Coutant et al., 2024), and can lead to malignant transformation into melanoma, an aggressive form of skin cancer responsible for most skin cancer-related deaths (Matthews et al., 2017).

The specification and differentiation of MCs from the neural lineage are governed by the melanocyte-inducing transcription factor (MITF). MITF regulates MC differentiation by controlling the expression of pigmentation genes (e.g., *Tyr*, *Trp1*, *Trp2*) and maintains cellular homeostasis by modulating genes involved, among others, in cell-cycle progression (e.g., *Cdk2*) and apoptosis (e.g., *Bcl2*) (Goding & Arnheiter, 2019; Kawakami & Fisher, 2017). MITF is also a major player of melanoma progression, influencing both, the melanoma cell differentiation state and plasticity, contributing to high tumor heterogeneity. Within a single tumor, different cell states co-exist: highly differentiated, proliferative melanocytic cells, which are associated with high MITF levels, whereas slow-cycling, dedifferentiated, invasive stem-like cells correlate with low MITF levels (Arozarena & Wellbrock, 2019; Carreira et al., 2006; Cheli et al., 2011; Hoek et al., 2008; Müller et al., 2014; Popovic & Tartare-Deckert, 2022; Rambow et al., 2019; Tirosh et al., 2016; Tsoi et al., 2018; Wouters et al., 2020). Interestingly, differentiated MCs can give rise to melanoma through a process that includes their reprograming and dedifferentiation (Köhler et al., 2017). However, the specific factors and mechanisms driving this dedifferentiation of mature MCs, and the role of MITF in these processes, remain poorly understood. Therefore, understanding how MITF affects MC plasticity and identifying novel factors that regulate MITF expression and activity could provide new insights into the mechanisms underlying MC transformation.

In addition to the heterotypic cell-cell interaction with keratinocytes, MCs are in direct contact with the basement membrane, a specialized extracellular matrix (ECM) rich in type IV collagen (COL IV), which forms the junction with the underlying dermis, comprising type I collagen (COL I) and fibronectin (FN) (Pfisterer et al., 2021). This implies that under physiological conditions, COL IV is the predominant ECM type epidermal MCs are exposed to, whereas interactions with dermal COL I and FN can occur due to altered basement membrane integrity, for instance, following injury or chronic UV exposure (Amano, 2009, 2016; Fisher & Rittié, 2018; Iriyama et al., 2011, 2020; Yoshihisa et al., 2014). Within the skin, a wide range of stiffness has been reported for different compartments, ranging from lower kPa (dermis-like) to lower MPa (epidermis-like) values (Biggs et al., 2020; Feng et al., 2022; Graham et al., 2019). Importantly, the cellular microenvironment, including ECM components and tissue stiffness, significantly impacts cell fate and function (Bonnans et al., 2014; Guilak et al., 2009; Walma & Yamada, 2020). Cells adapt to these molecular and mechanical parameters by transmitting environmental signals to intracellular signal transduction pathways, including mitogen-activated protein kinase (MAPK)/ extracellular signal-regulated kinase (ERK) pathways (Tan et al., 2023). Crucial for this process are focal adhesions (FAs), dynamic multi-protein complexes at the plasma membrane that link the ECM to the actin cytoskeleton via transmembrane receptors such as integrins. Notably, focal adhesion kinase (FAK), a key signaling protein within FAs, is activated through integrin engagement with the ECM, resulting in autophosphorylation and activation of downstream signaling pathways like MAPK/ERK (Paszek et al., 2005). In various cell systems, such signal activation downstream of ECM cues has been reported to affect the localization and activation levels of various transcription factors and coregulatory factors, such as Yes-associated protein (YAP), leading to gene expression changes that impact processes like proliferation, survival, or differentiation (Ishihara & Haga, 2022).

In melanoma cells, MAPK signaling components have recently been reported to negatively regulate MITF nuclear localization and activity in melanoma cells: rapidly accelerated fibrosarcoma (RAF) proteins, acting upstream of ERK, interact directly with MITF, causing its cytoplasmic retention and reduced transcriptional activity (Estrada et al., 2022), while MAPK/ERK signaling, in collaboration with glycogen synthase kinase 3 (GSK3), controls MITF nuclear export (Ngeow et al., 2018). The transcriptional co-activator YAP, known to translocate into the nucleus in response to ECM stiffening in various cell types (Cai et al., 2021; Dupont et al., 2011; Y. Huang et al., 2022; Miskolczi et al., 2018), promotes MITF expression in uveal and cutaneous melanoma cells (Barbosa et al., 2023; Miskolczi et al., 2018) and serves as MITF cofactor in uveal melanoma (Barbosa et al., 2023). This indicates a dual role of YAP in the control of MITF in melanoma. However, whether and how ECM cues modulate MITF remains to be elucidated.

So far, studies have separately explored the roles of ECM type and substrate stiffness for MC biology. For example, Hara et al. reported that COL IV stimulates dendricity in human MCs on glass substrates (considered as ultra-hard stiffness) (Hara et al., 1994), while melanin production on stiff PDMS substrates (5.5 MPa) coated with laminin is higher compared to softer counterparts (from 50 kPa to 1.8 MPa) (Choi et al., 2014). However, the potential synergistic effects of substrate stiffness and ECM components on MC functions remain to be defined. Understanding such effects is particularly important since the ECM undergoes continuous remodeling that includes de novo synthesis and degradation (Pfisterer et al., 2021). Various factors, such as skin aging, inflammation, sunlight exposure, wound healing, and fibrosis, can promote ECM remodeling, potentially resulting in aberrant ECM modifications (Pfisterer et al., 2021), which may modify the presentation of ECM ligands and alter substrate stiffness. For instance, during wound healing, the ECM transitions from a FN-rich provisional matrix to a COL I–rich structure as the tissue repairs. Nevertheless, it is still insufficiently understood how substrate stiffness influences the cellular responses to specific ECM components, and, conversely, whether cells need to engage in particular ECM interactions in order to sense environmental stiffness features.

This study investigated whether and how substrate stiffness and ECM components act together to modulate MC functions. We compared the combined effects of various ECM proteins and substrate stiffness on MC behavior. Given that MITF plays a central role in integrating environmental signals to regulate key aspects of MC identity, function, and plasticity—and considering its known regulation by the MAPK/ERK pathway, which mediates ECM signal transduction—we hypothesized that MITF may serve as a key effector of ECM-dependent cues in MCs. By investigating how ECM composition and mechanical properties affect MITF activity, we aimed to uncover mechanisms through which the microenvironment shapes MC behavior and plasticity. Our results revealed that ECM components control MC differentiation and function via a FAK-MEK-ERK-MITF axis, with different ECM types determining the ability of MCs to perceive and respond to mechanical stimuli in their environment.

## RESULTS

### ECM components differentially affect MC morphology, adhesion and migration

To assess how extracellular mechanical cues affect MC behavior, we employed PDMS substrates of varying stiffnesses: ≈45 kPa defined here as soft (fig. S1A), ≈140 kPa as intermediate (fig. S1B) and ≈900 kPa as stiff (fig. S1C). This range was chosen to reflect the wide range of stiffness found within the skin and because increased environmental stiffness is associated with multiple pathologies (Diazzi et al., 2020; Pfisterer et al., 2021; K. Wang et al., 2023). To mimic both normal and altered ECM conditions, COL IV-, COL I-, and FN-coated substrates of varying stiffness were used for cultures of either primary (pMCs) or immortalized murine MCs (stably expressing membrane-targeted tandem dimer Tomato; iMCs, generated for this study), followed by molecular, cellular and functional assays.

Interestingly, visual inspection in phase-contrast and fluorescence microscopy revealed distinct ECM-specific phenotypes, with COL I and FN eliciting opposing cellular responses. iMCs cultured on FN-coated substrates exhibited a more dendritic shape compared to the mostly bi- or tripolar MCs grown on COL I (fig. 1A). To examine this phenotype further, we performed a Sholl analysis (fig. 1A-C; fig. S2A; Binley et al., 2014; Sholl, 1953). Quantification of the dendritic complexity of MCs (fig. 1B) and the corresponding Sholl profile (fig. 1C; fig. S2A) confirmed that most iMCs exposed to COL IV and FN displayed between 4 to 7 dendrites, whereas MCs grown on COL I exhibited mostly 2 to 3 dendrites (fig. 1B). Noteworthy, over the range of stiffnesses tested, MC dendricity was not impacted by substrate stiffness, irrespective of the ECM protein type used (fig. S2A).

**Figure 1:**
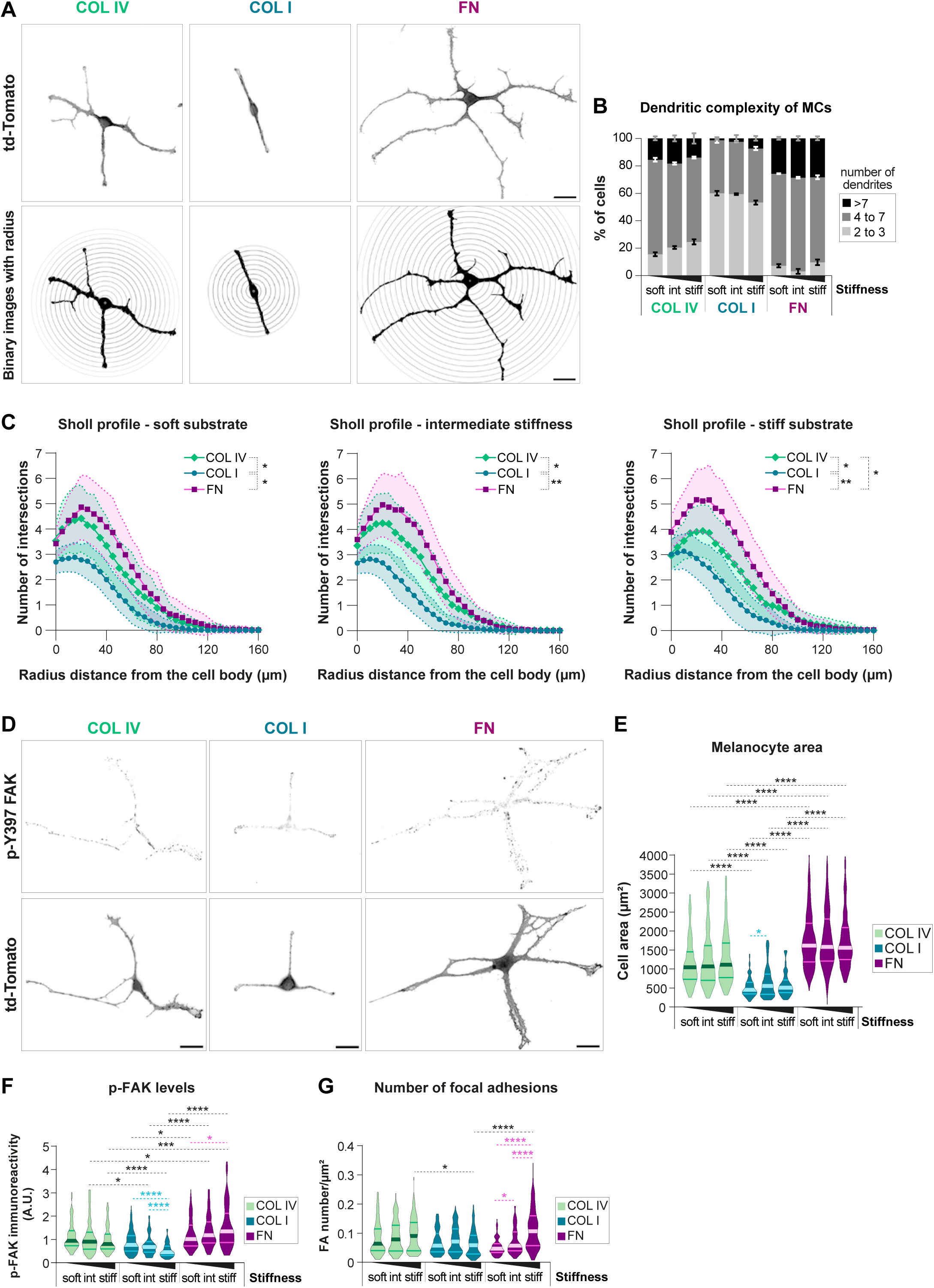
ECM components determine the morphology and adhesion of immortalized melanocytes. iMCs were cultured overnight on substrates of varying stiffness defined here as soft (PDMS ratio 1:60 ≈40kPa), intermediate (PDMS ratio 1:35 ≈130kPa) and stiff (PDMS ratio 1:10 ≈1MPa) coated with COL IV, COL I or FN. **A. Top:** Representative micrographs of iMCs, expressing td-Tomato, cultured overnight on stiff substrates coated with COL IV, COL I or FN; scale bar: 20µm. **Bottom**: Representative binary image of cells used for Sholl analysis, with radius drawn every 5µm. **B.** Quantification of the number of dendrites of iMCs; graph indicating the percentage of cells exhibiting 2 or 3 dendrites, between 4 and 7, or more than 7 dendrites; mean±SEM, N=3, n(cells)≥85. **C.** Graphs showing the Sholl profile analysis, plotting the number of dendrite intersections against the distance from the cell body. The SEM are represented by the connecting curve (dotted line); N=3, n(cells)≥85. Each curve represents an ECM type, and each graph represents a substrate stiffness condition; Welch ANOVA test: **: p<0.005; *: p<0.05. **D.** Representative micrographs of iMCs (expressing membrane-targeted tandem dimer (td) Tomato) cultured on stiff substrates coated with COL IV, COL I or FN and immunostained for p-Y397-FAK and with DAPI; scale bar: 20µm. **E.** Quantification of the cell area of iMCs; violin plots display the medians and distributions of cell area in each condition; N=3, n(cells)≥69; Kruskal–Wallis test: ****: p<0.0001; **: p<0.005; *: p<0.05. **F.** Quantification of p-FAK levels; violin plots show the medians and distributions of the integrated density of p-FAK per cell; N≥4, n(cells)≥82; Kruskal–Wallis test: ****: p<0.0001; ***: p<0.001; *: p<0.05. **G.** Quantification of the number of focal adhesions per µm^2^ per cell with violin plots displaying medians and distributions; N≥3, n(cells)≥65; Kruskal– Wallis test: ****: p<0.0001; *: p<0.05. Abbreviations: A.U., arbitrary units.

Since cell spreading is regulated by coordinated changes in adhesions to ECM and cytoskeletal reorganization, we next examined the cell area as well as key cell adhesion parameters. Among these, FAK activity by means of its autophosphorylation (p-FAK) and quantification of the number of focal adhesions (FAs) per cell served as indicators of responses resulting from MC-ECM interactions. In both pMCs and iMCs, the smallest cell area was detected on COL I (fig. 1D,E; fig. S3A) and correlated with the lowest p-FAK levels (fig. 1D,F; fig. S3B), whereas the largest cell areas (fig. 1D,E; fig. S3A) were found on FN, correlating with high p-FAK signals (fig. 1D,F; fig. S3B). Substrate stiffness, however, seemed to have less influence on the cell area: apart from a slight increase from soft to intermediate stiffness in iMCs cultured on COL I, we did not note significant changes of the cell area across the different substrate stiffnesses, regardless of the ECM protein used (fig. 1E; fig. S3A). In contrast, both p-FAK and the number of FAs per µm² exhibited a significant stiffness-mediated increase in MCs grown on FN-coated substrates (fig. 1F,G; fig. S3B,C). In contrast, the overall low p-FAK levels in COL I-exposed iMCs even decreased with stiffness (fig. 1F), highlighting that stiffness sensing in MCs depends on the ECM type.

Finally, since cell morphology and FAs are closely linked to cell movement, we investigated whether the distinct ECM molecules influence MC motility. Live cell imaging of iMCs revealed that both the distance traveled (fig. 2A-C) and velocity (fig. 2D) of cell migration decreased on COL I compared to COL IV and FN, whereas FN-grown iMCs showed increased motility (fig. 2A-C) and faster migration (fig. 2D) relative to their COL IV-grown counterparts.

**Figure 2:**
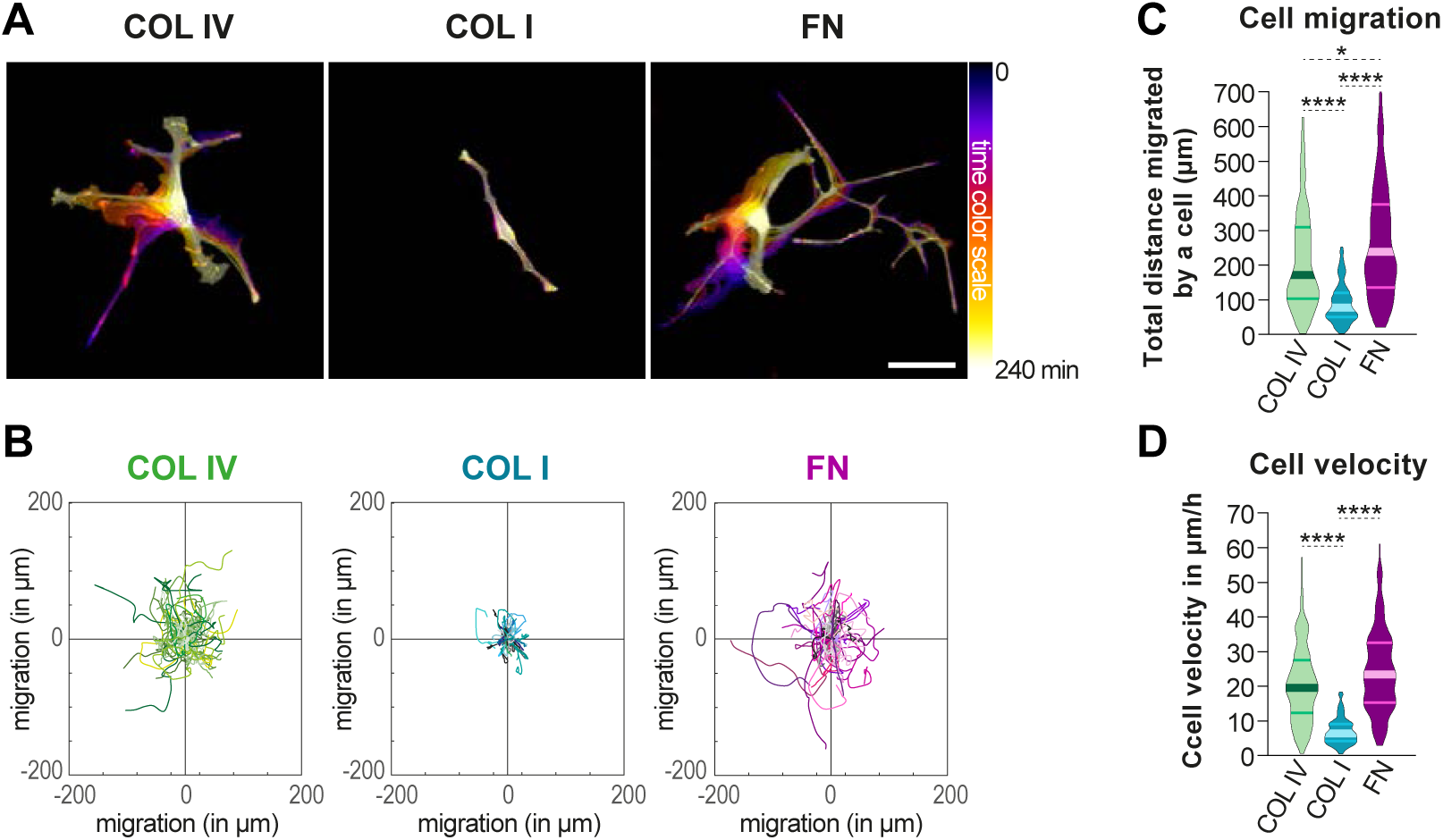
Differential effects of ECM proteins on iMC motility, with COL I decreasing and FN increasing motility. **A.** Image showing the temporal color code analysis from live-cell imaging of iMCs cultured overnight in plastic chamber coated with COL IV, COL I or FN to visualize cell dynamics over time. The different colors represent morphology and position of the iMCs at a given time (20min intervals), with the color gradient indicating time progression (from 0 to 240min); scale bar: 50µm. **B**. Tracks showing the path and distance of cells (one cell = one track) relative to the point of origin (time point zero) in the x and y plane. **C**. Quantification of total distance traveled by iMCs; violin plots show the medians and distributions; N=3, n(cells)≥113; Kruskal–Wallis test: ****: p<0.0001; *: p<0.05. **D**. Quantification of mean speed of iMCs; violin plots show the medians and distributions N=3, n(cells)≥113; Kruskal–Wallis test: ****: p<0.0001.

Together, these data show that COL I and FN trigger opposite morphological, adhesive and migratory phenotypes in MCs.

### COL I and FN have opposite effects on melanin production and MC proliferation

Given the observed role of ECM components for MC morphology and migration, we next investigated whether ECM cues influence other MC functions. As a hallmark task of MCs, we examined their melanin production by measuring both intracellular melanin contents and melanin released into the culture medium. iMC grown on COL I-coated substrates exhibited the highest intra- and extracellular melanin content, while melanin levels on FN were lowest compared to COL I and COL IV (fig. 3A,B). Notably, the expression of *Tyr*, an essential gene for melanogenesis, correlated with melanin production, exhibiting a higher expression in iMCs exposed to COL I than in those exposed to FN (fig. 3C). Like iMCs, pMCs cultured on FN produced least melanin on soft and intermediate stiffness (fig. S3D). For all ECM types tested, highest values for melanin production were observed at high stiffness (fig. S3D, fig. 3A,B). Collectively, these data demonstrated that compared to COL I, FN restricts melanin production.

**Figure 3:**
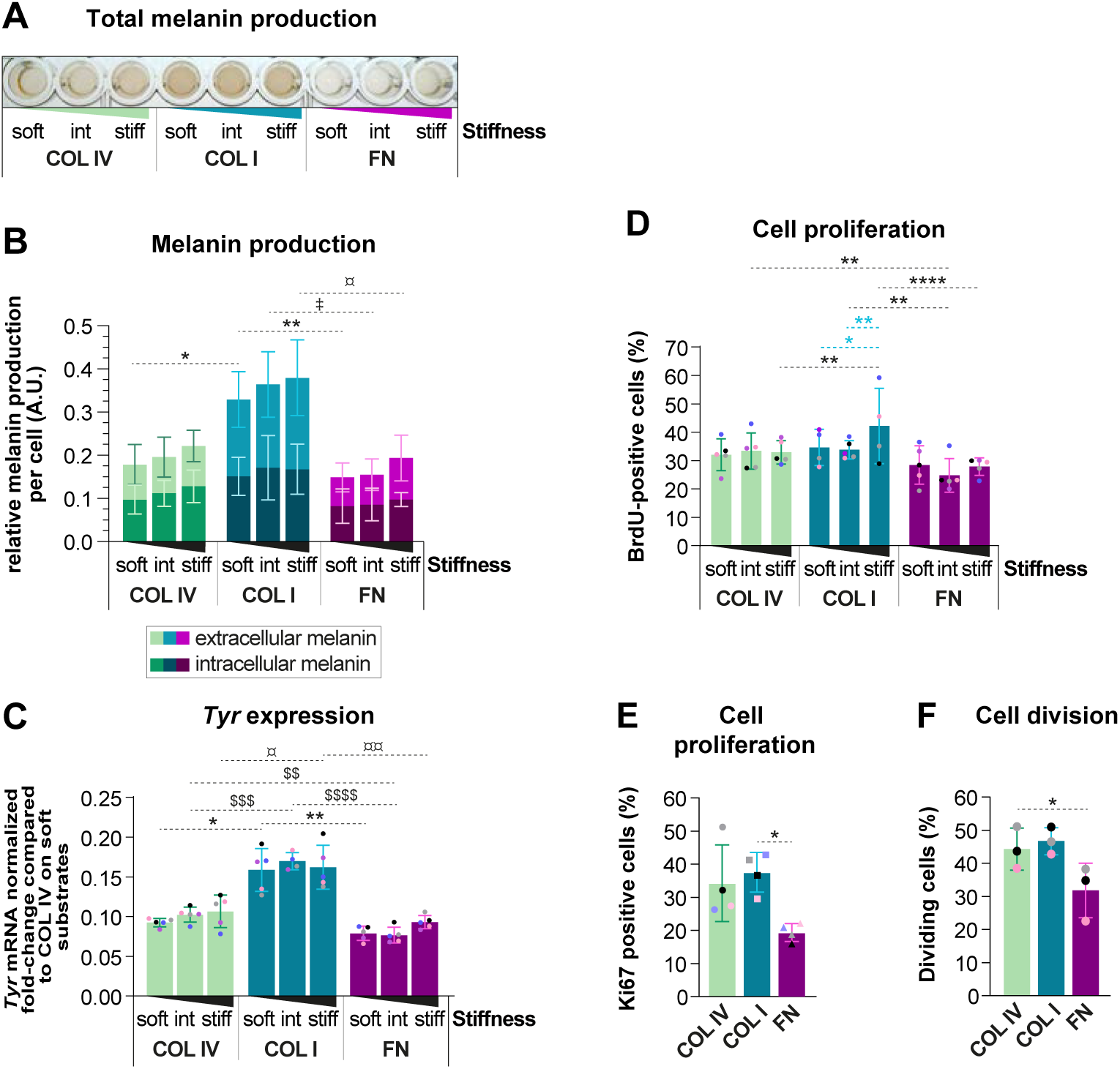
COL I stimulates melanin production, and FN counteracts iMC proliferation. **A**. Representative example of total melanin produced by iMCs cultured 72 h on substrates of varying stiffness (soft, intermediate or stiff) coated with COL IV, COL I or FN. **B.** Quantification of intra- and extracellular melanin content by spectrophotometry at 405 nm from iMCs cultured as in (A); means±SD, N≥5; Welch ANOVA test: **: p<0.005; *: p<0.05. **C.** Quantification of *Tyr* mRNA expression of iMCs cultured overnight on substrates of varying stiffness (soft, intermediate or stiff) coated with COL IV, COL I or FN; means±SD, N=5; Kruskal–Wallis test: **: p<0.005; *: p<0.05. **D**. Percentage of BrdU-positive cells from iMCs cultured 48 h on substrates of varying stiffness (soft, intermediate or stiff) coated with COL IV, COL I or FN; means±SD, N≥4; ordinary two-way ANOVA: ****: p<0.0001; ***: p<0.001; **: p<0.005; *: p<0.05. **E**. Percentage of Ki67-positive iMCs cultured 48 h on stiff substrates; means±SD, N=4; Welch ANOVA test: *: *p* < 0.05. **F**. Percentage of dividing cells from live-cell imaging of iMCs cultured overnight on COL IV, COL I and FN; means±SD, N=3; RM one-way ANOVA test: *: p<0.05. Abbreviations: A.U., arbitrary units.

Using a BrdU assay, we next evaluated MC proliferation. While stiffness had no effect on MC proliferation in the presence of COL IV and FN, we observed a higher proportion of BrdU-positive iMCs when exposed to stiff COL I-coated substrates, indicating a synergistic effect of COL I and stiffness on MC proliferation (fig. 3D). Among the three ECM types tested, FN exposure resulted in the lowest proliferation rates (fig. 3D), a finding that could be further confirmed by Ki67 immunostaining (fig. 3E), as well as live cell imaging followed by quantification of the percentage of dividing cells (fig. 3F). Overall, these experiments revealed a stiffness-dependent proliferative phenotype triggered by COL I, while FN appears to restrict cell division of MCs.

Together, our findings show that melanin production and *Tyr* expression are highest in MCs exposed to COL I and lowest in FN-grown MCs. Moreover, cell proliferation was lowest on FN, and stiffness-mediated increase in proliferation was only observed on COL I-coated substrates, suggesting that MC mechanoresponses depend on the ECM type.

### MITF nuclear localization is modulated by ECM molecules and substrate stiffness but does not correlate with nuclear YAP

Our data indicate an ECM-dependent regulation of MC proliferation and melanin production, with COL I and FN driving opposite effects on these cellular functions. Considering the central role of the transcription factor MITF in the control of both melanogenesis and cell cycle progression (Goding & Arnheiter, 2019), and taking into account the observed increase of the MITF target gene *Tyr* on COL I (fig. 3C), we next examined the impact of ECM cues on the protein levels and subcellular localization of MITF (fig. 4A-C; fig. S3E). Although overall MITF levels were lowest on COL I, nuclear MITF levels were highest on this ECM type (fig. 4A-C), consistent with pronounced melanin production. In contrast, MCs cultured on FN displayed low nuclear MITF levels compared to COL I-coated substrates (fig. 4A-C; fig. S3E).

**Figure 4:**
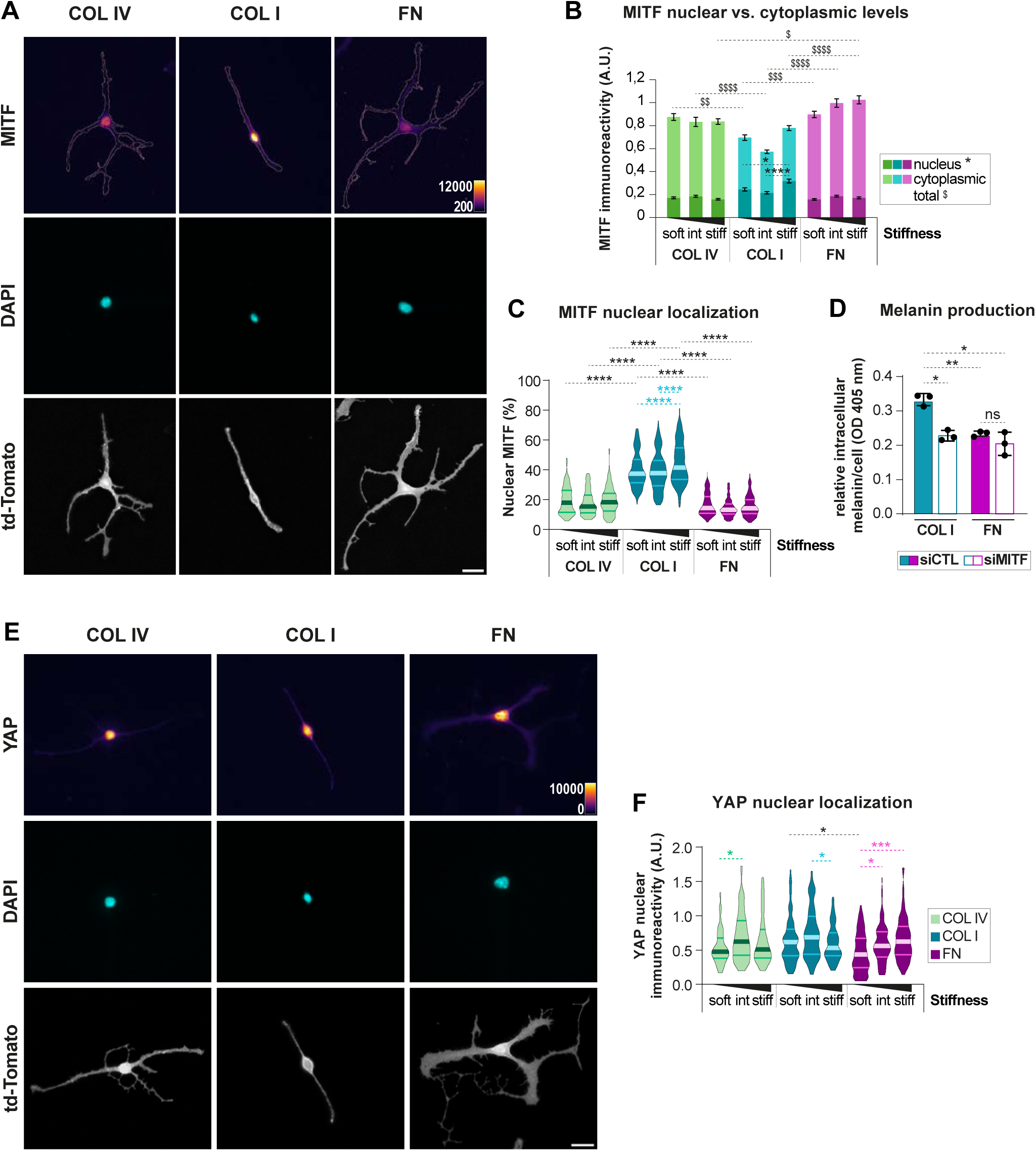
MITF nuclear localization is modulated by both ECM molecules and substrate stiffness. **A.** Representative micrographs of iMCs expressing td-Tomato cultured overnight on stiff substrates coated with COL IV, COL I or FN and stained for MITF and DAPI; scale bar: 20µm. A pseudo-color intensity scale (’fire’ color map) was applied to enhance visualization of MITF nuclear localization. **B**. Quantification of cytoplasmic and nuclear MITF levels; graphs show the mean±SEM of nuclear and cytoplasmic MITF intensities per cell 24 h post-plating on soft, intermediate, or stiff substrates coated with COL IV, COL I, or FN; N≥3, n(cells)≥89; Kruskal–Wallis test for nuclear MITF: ****: p < 0.0001; *: p<0.05; Kruskal–Wallis test for total MITF: $$$$: p < 0.0001; $$$: p < 0.001; $$: p<0.005; $: p<0.05.**C**. Quantification of nuclear MITF levels; violin plots show the medians and distributions of the percentage of nuclear MITF per cell; N≥3, n(cells)≥67; Kruskal–Wallis test: ****: p<0.0001; *: p<0.05. **D**. Quantification of intracellular melanin levels following siRNA-mediated MITF knockdown. iMCs were transfected with control siRNA (siCtrl) or MITF-targeting siRNA (siMITF) 4 h after plating on COL I or FN. Seventy-two hours post-transfection, cells were harvested and melanin content was measured by spectrophotometry at 405 nm; means±SD, N=3; Welch ANOVA test: **: p<0.005; *: p<0.05. **E**. Representative micrographs of iMCs expressing td-Tomato cultured overnight on stiff substrates coated with COL IV, COL I or FN and stained for YAP and DAPI; scale bar: 20µm. A pseudo-color intensity scale (’fire’ color map) was applied to enhance visualization of YAP nuclear localization. **F**. Quantification of nuclear YAP levels; violin plots show the medians and distributions of the nuclear integrated density of YAP per cell; N=3, n(cells)≥71; Kruskal–Wallis test: ***: p<0.001; *: p<0.05. Abbreviations: A.U., arbitrary units; td-Tomato, membrane-targeted tandem dimer Tomato.

Furthermore, a stiffness-mediated increase of MITF nuclear localization was noted in MCs on COL I (fig. 4B,C; fig. S3E), thus identifying MITF as an ECM- and mechanosensitive transcription factor. The contrasting phenotypes of MCs grown on COL I vs. FN were also evident when correlating nuclear MITF levels to the cell area (fig. S4A): while MCs grown on COL I were characterized by a low ratio of cell area to nuclear MITF compared to COL IV, FN-exposed MCs showed high ratios, i.e. a large cell area was associated with low MITF. Interestingly, this ratio decreased with substrate stiffness on COL I and FN but not on COL IV (fig. S4A), suggesting that changes in nuclear MITF are not linearly related to morphological changes in response to ECM. siRNA-mediated knock-down of MITF (fig. S4B) confirmed the principal role of MITF for melanin production on both COL I and FN (fig. 4D).

Given that YAP is a transcriptional co-activator known to translocate into the nucleus in response to ECM stiffening in various cell types, including melanoma cells (Miskolczi et al., 2018), and that YAP may serve as potential MITF partner in uveal melanoma cells (Barbosa et al., 2023), we wondered whether YAP could be associated with the observed ECM-dependent MC phenotypes. We examined YAP subcellular localization and, consistent with previous reports, observed a stiffness-mediated increase in nuclear YAP, with the strongest response at intermediate stiffness levels on COL IV and COL I, and on stiff FN-coated substrates (fig. 4E,F). Importantly, however, unlike nuclear MITF, we hardly observed significant differences of nuclear YAP when comparing the three ECM types, making it unlikely that MCs utilize YAP for ECM-specific melanin production.

Collectively, our findings suggest that ECM subtypes regulate MC behavior and function likely through modulation of MITF localization and activity, independently of YAP.

### Bulk RNA sequencing confirms that ECM types elicit significant changes in expression of genes associated with MC pigmentation and differentiation

Our data so far indicate that ECM composition modulates MITF localization and activity in MCs, suggesting that ECM components may influence MITF-regulated transcriptional programs. To investigate this further and gain a broader understanding of how ECM components shape the MC phenotype, we performed bulk RNA sequencing (RNAseq) on MCs cultured on COL IV, COL I, and FN (fig. 5; fig. S5). To capture global transcriptomic changes associated with distinct ECM environments and to identify gene signatures linked to MC differentiation, plasticity, and other relevant pathways, we analyzed the expression of MITF target genes, MC-specific genes, and broader signaling networks that are associated with the phenotypic shifts observed across ECM conditions.

**Figure 5:**
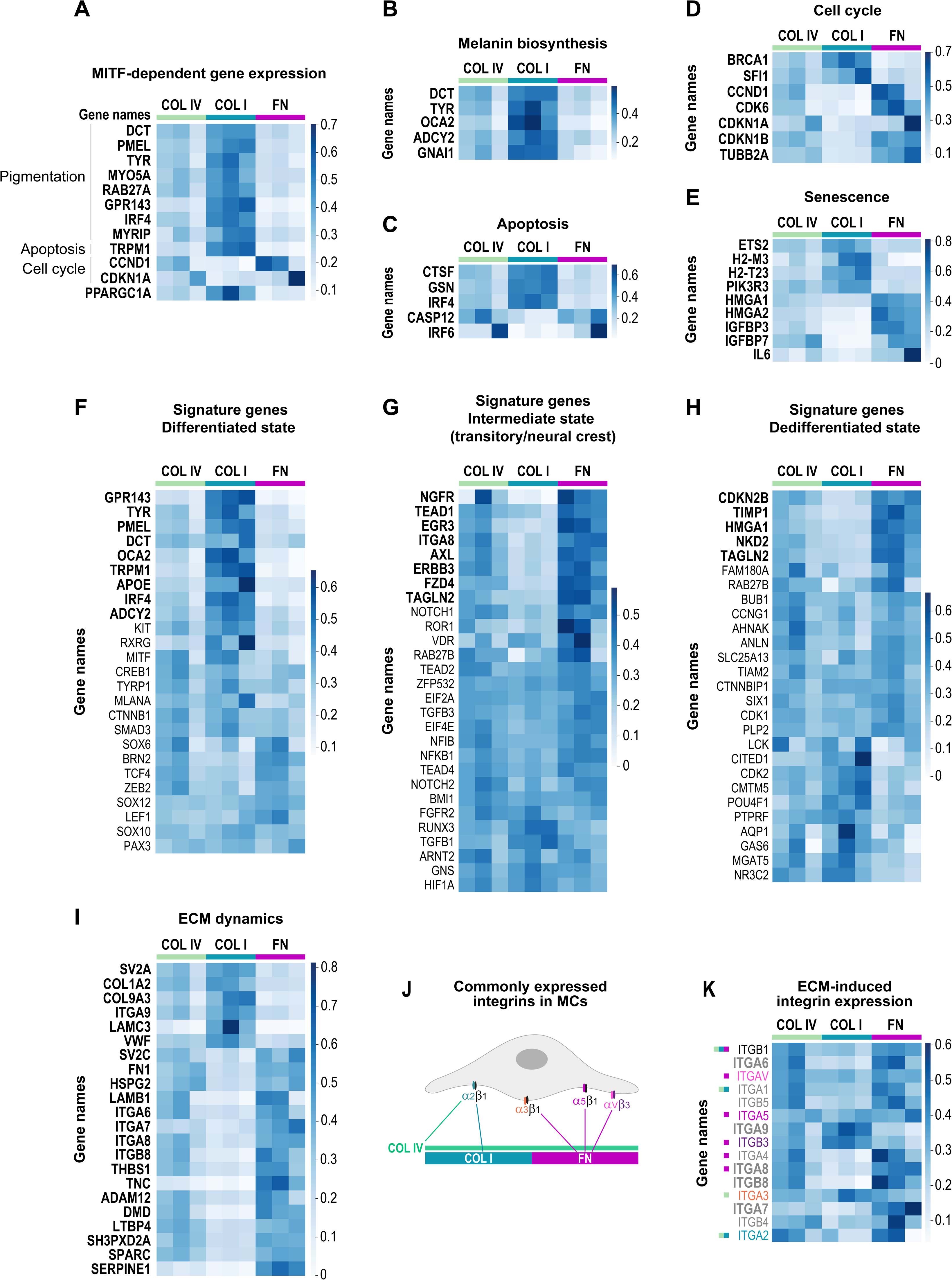
ECM-dependent transcriptional changes in iMCs. iMCs were cultured overnight on stiff substrates coated with COL IV, COL I or FN. Gene expression heatmaps showing transcript abundance across ECM conditions. Darker blue indicates higher expression; lighter blue indicates lower expression; bold labels indicate statistically differentially expressed genes. Displayed values represent normalized expression levels per gene across conditions (e.g., standard scaling), enabling comparison of the same gene between ECM conditions despite differences in expression range. Panels show: **A.** differentially expressed MITF-dependent genes; **B.** genes involved in melanin biosynthesis; **C–E.** genes related to apoptosis, cell cycle, and senescence, respectively; **F–H.** signature genes of the differentiated, intermediate, and dedifferentiated states; genes were grouped by clustering of expression profiles; **I.** genes involved in ECM dynamics; **J.** schematic overview of integrins commonly expressed in MCs; and **K.** integrin gene expression across COL IV, COL I, and FN conditions.

Comparison of global transcriptomic profiles across the ECM types used revealed a range of differentially expressed genes (DEGs) (fig. 5; fig. S5B,C). Consistent with the reduced melanin production observed for MCs on FN (fig. 3B), several MITF-dependent target genes, which are also pigmentation-related (e.g., *Dct*, *Pmel*, *Tyr*, *Gpr143*), were significantly downregulated on FN compared to COL I (fig. 5A,B). In addition, further components involved in melanin biosynthesis and trafficking—such as *Oca2*, *Adcy2*, and *Rab27a*—also showed reduced expression on FN (fig. 5A,B). Of note, some of the MITF-dependent DEGs have previously been linked to apoptosis *(Trpm1)* and cell cycle progression (*Ccnd1*, *Cdkn1a*) (fig. 5A). Global gene expression analysis indeed revealed further DEGs related to cell cycle control and apoptosis (fig. 5C,D), in line with the observed decrease in proliferation of MCs cultured on FN when compared to COL I (fig. 3D,E). In addition, FN elicited differential expression of genes associated with cellular senescence (fig. 5E). Taken together, these findings suggest that adhesion to FN promotes a transcriptional program in MCs that may counteract cell cycle progression.

Considering that MITF-dependent gene expression overlapped with transcriptional programs involved in melanocyte development, and that these programs were downregulated in MCs cultured on FN (fig. 5A), we extended our analysis to gene expression signatures associated with melanoma plasticity and differentiation states (Arozarena & Wellbrock, 2019; Durand et al., 2024; F. Huang et al., 2021; Konieczkowski et al., 2014; Pessoa et al., 2021; Rambow et al., 2015, 2019; Tsoi et al., 2018). Comparing FN to COL I revealed downregulation of genes associated with melanocytic differentiation on FN, such as *Apoe,* next to the previously mentioned MITF target genes (e.g., *Dct*, *Pmel*, *Tyr*, *Trpm1*, *Gpr143*, *Irf4*) and melanin biosynthesis-related genes (e.g., *Oca2*, *Adcy2)* (fig. 5A,B,F). However, core melanocytic factors such as *Sox10*, *Lef1*, *Creb1*, *Pax3*, and *Mlana* (fig. 5F) were not significantly altered between COL I and FN, suggesting maintenance of lineage identity. In contrast, several genes associated with cell plasticity *(Tead1, Fzd4*, *Tagln2)* and neural crest-like features (*Ngfr, Erbb3*) were significantly upregulated on FN relative to COL I (fig. 5G). Additionally, genes previously associated with a dedifferentiated state, such as *Axl, Itga8*, *Timp1*, *Hmga1*, and *Nkd2*, were upregulated on FN compared to COL I (fig. 5H), further supporting the emergence of a phenotype characterized by increased plasticity and reduced differentiation on FN.

Gene set enrichment analysis (GSEA) identified “ECM organization” as the top enriched Reactome pathway in iMCs cultured on FN compared to COL I (fig. S5E), indicating transcriptional remodeling of ECM-associated genes in response to ECM molecules. Among these, collagen genes (*Col1a2* and *Col9a3;* fig. 5I) were downregulated on FN, while non-collagenous ECM components, including *Fn1*, *Tnc*, *Hspg2* and *Thbs1,* showed increased expression (fig. 5I). In addition, ECM-modifying genes (*Sparc*, *Adam12*, *Serpine1;* Fig. 5I) were also upregulated, supporting a potential active ECM remodeling on FN. Together, these gene expression patterns align with the emergence of a cell state characterized by increased migratory potential and plasticity in MCs grown on FN.

Interestingly, several integrin subunits were also differentially expressed across ECM conditions (fig. 5I). A broad repertoire of integrins was found to be expressed in iMCs grown on COL IV, COL I, and FN, including subunits commonly reported in MCs (Arias-Mejias et al., 2020; Hara et al., 1994; Morelli et al., 1993; Pinon & Wehrle-Haller, 2011; Scott et al., 1992), such as *Itga2, Itga3, Itga5, ItgaV, Itgb1,* and *Itgb3*, which mediate adhesion to collagens and fibronectin (fig. 5J, K). While most of these were expressed independently of the ECM type used, we also noted that *Itga6*, *Itga7*, *Itga8*, and *Itgb8* were upregulated on FN, while *Itga9* was upregulated on COL I (fig. 5I). Taken together, our data suggests that while ECM composition influences integrin gene expression to some extent, changes in integrin profiles alone may not fully account for the phenotypic differences observed between the distinct ECM components.

### ECM-dependent FAK/MEK/ERK activation controls MITF localization and activity, as well as melanogenesis

Given the ECM-dependent phenotypic shifts in MCs, we next analyzed expression changes in key signaling pathways— PI3K/Akt (Larribere et al., 2004; Phung et al., 2011; Shi et al., 2016; C. Wang et al., 2016), Wnt (Colombo et al., 2022; Katkat et al., 2023; Sinnberg et al., 2018), and MAPK (Buscà & Ballotti, 2000; Estrada et al., 2022; Ngeow et al., 2018; Ostojić et al., 2021; Wellbrock & Arozarena, 2015)— known to regulate MITF activity, melanocyte differentiation, and plasticity. While only a few PI3K/Akt-associated genes were differentially expressed, with *Brca1* and *Sgk2* downregulated and *Sgk1* upregulated on FN (fig. 6A), Wnt signaling showed a broader modulation (fig. 6B). Among the DEGs, multiple components— including the canonical target *Ccnd1*, the positive regulators *Frat2*, *Fzd4*, and *Cd88c*, and the feedback inhibitor *Nkd2*—were upregulated on FN, suggesting an activation of both canonical and non-canonical Wnt signaling pathways, potentially accompanied by feedback regulation (fig. 6B). The most pronounced transcriptional changes, however, were observed in MAPK (fig. 6C) and ERK signaling (fig. 6D), reflected by a high number of DEGs associated with these pathways. Some upstream activators of ERK (*Cnksr1*, *Rasgrp3*, *Pak3*) were downregulated on FN compared to COL I (fig. 6C). In contrast, genes enhancing ERK signaling (*Igf2bp1*, *Irak2*, *Il1rap*, *Ngfr*) and transcriptional ERK targets (*Etv4*, *Il6*, *Tnc*, *Lif*, *Col1a2*) were upregulated on FN-coated substrates (fig. 6C, D). In addition, upstream modulators of ERK, such as *Erbb3*, *Axl*, and *Itga6*, were also upregulated (fig. 6C, D).

**Figure 6:**
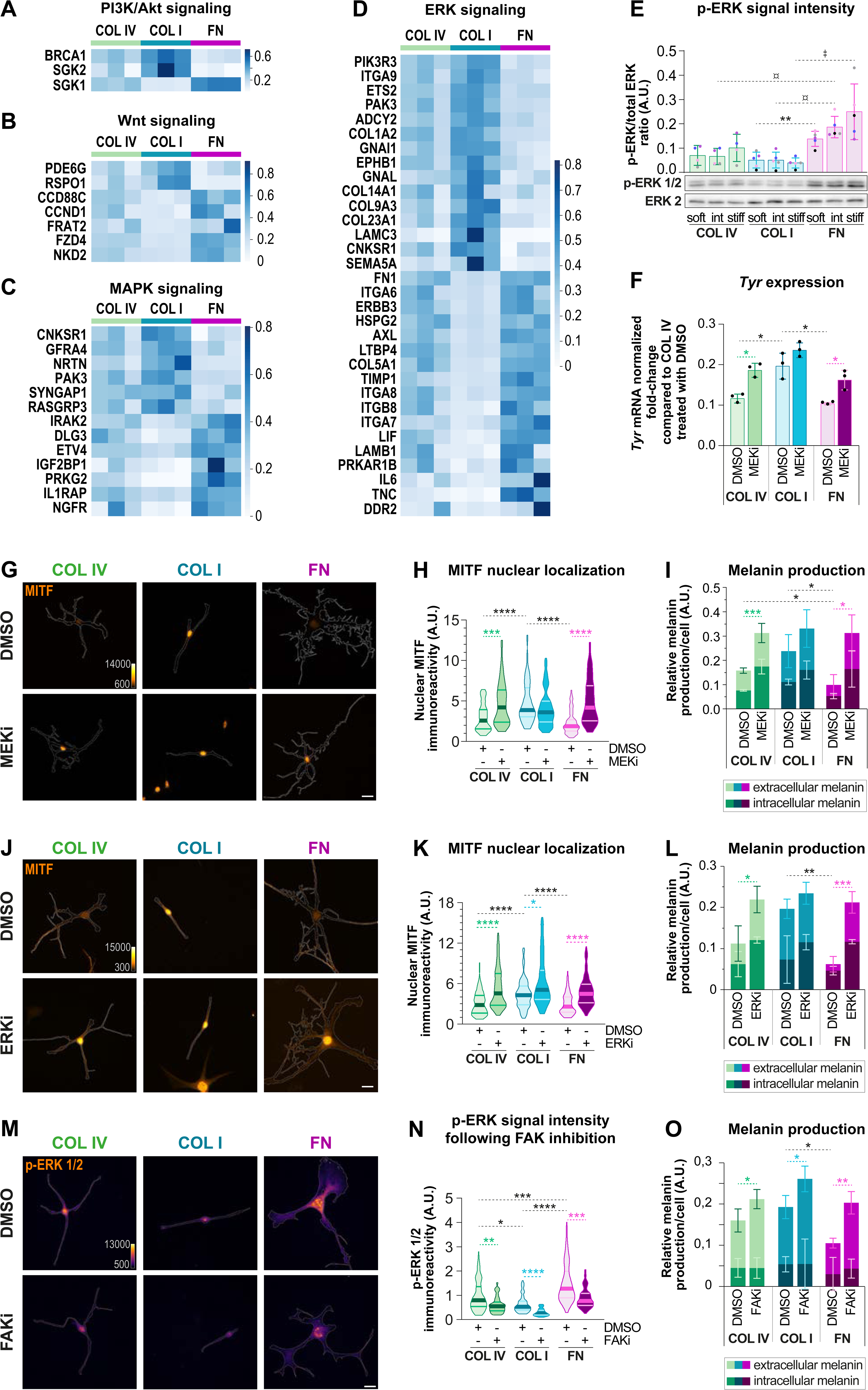
ECM-mediated ERK activation regulates MITF localization and melanogenesis. **A-D.** iMCs were cultured overnight on stiff substrates coated with COL IV, COL I or FN. Gene expression heatmaps of signaling pathway components across ECM conditions. Darker blue indicates higher transcript abundance; lighter blue indicates lower abundance; bold labels denote statistically differentially expressed genes. Heatmaps illustrate: **A.** genes involved in the PI3K/Akt signaling; **B.** components of the Wnt signaling; **C.** genes associated with MAPK signaling; and **D.** genes associated with MAPK signaling. **E.** iMCs were cultured overnight on substrates of varying stiffness (soft, intermediate or stiff) coated with COL IV, COL I or FN. Representative Western blot and associated graph depicting the quantification of p-ERK1/2 Thr202/Tyr204 to total ERK2 ratio; means±SD; COL IV, N=4; COL I and FN, N=5; Welch ANOVA test: **: p<0.005 and ‡ : p<0.05; and Kruskal– Wallis test: ¤ : p<0.05. **F-O.** iMCs were cultured on stiff substrates coated with COL IV, COL I or FN. 4 h post-plating, a 16-hrs (F-H) or 72-hrs (I) treatment with DMSO or 100 nM of MEK inhibitor (MEKi = Trametinib), a 16-hrs (J, K) or 72-hrs (L) treatment with DMSO or 10 µM of ERK inhibitor (ERKi = Ravoxertinib) or a 16-hrs (M, N) or 72-hrs (O) treatment with DMSO or 10 µM of FAK inhibitor (FAKi = Ifebemtinib) was commenced. **F.** Quantification of *Tyr* mRNA expression after MEK inhibitor treatment; means±SD, N=3; multiple t tests: **: p<0.005; *: p<0.05. **G**. Representative immunostainings for MITF following MEK inhibition; scale bar: 10µm. A pseudo-color intensity scale (“orange hot” color map) was applied to enhance visualization of MITF nuclear localization. **H**. Quantification of (G); violin plots showing the medians and distributions of the integrated density of nuclear MITF per cell; N=3, n(cells)≥63; Kruskal–Wallis test: ****: p<0.0001; ***: p<0.001. **I**. Quantification of intra- and extracellular melanin content following MEK inhibition by spectrophotometry at 405 nm; means±SD, N=3; multiple t tests: ***: p<0.001; *: p<0.05. **J**. Representative immunostainings for MITF following ERK inhibition; scale bar: 20µm. A pseudo-color intensity scale (“orange hot” color map) was applied to enhance visualization of MITF nuclear localization. **K**. Quantification of (J); violin plots showing the medians and distributions of the integrated density of nuclear MITF per cell; N=3, n(cells)≥89; Kruskal–Wallis test: ****: p<0.0001; *: p<0.05. **L**. Quantification of intra- and extracellular melanin content following ERK inhibition by spectrophotometry at 405 nm; means±SD, N=4; Welch ANOVA tests: ***: p<0.001; **: p<0.005; *: p<0.05. **M**. Representative immunostainings for p-ERK following FAK inhibition; scale bar: 20µm. A pseudo-color intensity scale (“fire” color map) was applied to enhance visualization of p-ERK intensity. **N**. Quantification of (M); violin plots showing the medians and distributions of the integrated density of p-ERK per cell; N=3, n(cells)≥83; Kruskal–Wallis test: ****: p<0.0001; ***: p<0.001; **: p<0.005; *: p<0.05. **O**. Quantification of intra- and extracellular melanin content following FAK inhibition by spectrophotometry at 405 nm; means±SD, N=5; multiple t tests: ***: p<0.001; **: p<0.005; *: p<0.05. Abbreviations: A.U., arbitrary units.

These data pointed to ECM-driven remodeling of MAPK/ERK signaling, prompting us to directly assess ERK activation in iMCs grown on COL IV, COL I, and FN. Immunoblot analyses of ERK1/2 phosphorylation (p-ERK) revealed an ECM-dependent modulation of ERK activity, with FN-grown iMCs featuring highest p-ERK levels, and COL I-exposed MCs displaying lowest p-ERK levels (fig. 6E). Considering these differential p-ERK signals observed with the various ECM types and given that in melanoma cells, MAPK/ERK has recently been shown to negatively control MITF nuclear localization and activity (Estrada et al., 2022; Ngeow et al., 2018), we next asked whether MEK/ERK activation was causal for the observed ECM-dependent MC responses. For this, we used Trametinib, a pharmacological inhibitor targeting MEK, the upstream kinase mediating ERK phosphorylation and activation (Gilmartin et al., 2011). Next to the expected loss of ERK phosphorylation (fig. S6A), Trametinib treatment (indicated as MEKi) increased *Tyr* mRNA expression (fig. 6F) and resulted in increased nuclear MITF in iMCs exposed to COL IV and FN, reaching the nuclear MITF levels observed on COL I (fig. 6G,H). Congruent with this, melanin production by iMCs cultured on both COL IV- and FN-coated substrates and treated with Trametinib were significantly elevated towards those levels observed in COL I conditions (fig. 6I). Similar results were obtained using an ERK inhibitor Ravoxertinib (Blake et al., 2016; Varga et al., 2020) (fig. 6J-L). This indicates that MEK/ERK activity in MCs negatively regulates nuclear MITF levels. We next aimed to delineate signals that might transduce ECM sensing to activation of ERK. Given the correlation between elevated FAK and ERK activity in FN-exposed MCs and considering that FAK has been reported to transduce ECM signals through MAPK/ERK (Paszek et al., 2005), we inhibited FAK in MCs. As expected, treatment of MCs with the FAK inhibitor Ifebemtinib (Li et al., 2021) resulted in reduced p-FAK levels as well as a reduced number of FAs in MCs (fig. S6B, C). More importantly, FAK inhibition resulted in a reduction of p-ERK levels as assessed by immunocytochemistry (fig. 6M,N) and caused a significant increase in melanin production (fig. 6O). Together, these data demonstrate that FN-mediated MEK/ERK activation counteracts MITF nuclear accumulation and melanin synthesis, suggesting that MEK/ERK — potentially downstream of FAK— act as negative regulators of melanogenesis in response to ECM cues.

In summary, we deciphered that distinct ECM proteins in the MC environment steer melanogenesis through differential activation of the MEK/ERK pathway and subsequent regulation of MITF localization and activity. We further showed that COL I and FN exert opposite effects on MC behavior and functions: COL I elicits a highly pigmented and proliferative, but non-motile MC state, while FN triggers a less pigmented, low proliferative and highly motile MC state. Finally, by combining varying substrate stiffness and distinct ECM types we delineated that stiffness-mediated MC functions rely on specific ECM components.

## DISCUSSION

### ECM components fine-tune MC phenotypes and functions

Our study identifies ECM components as critical environmental triggers that instruct MC behavior. Through dynamic interactions with the ECM, MCs engage adhesion-dependent signaling, such as FAK activation, enabling them to decode contextual ECM inputs and adapt their phenotype accordingly. Specifically, we observed that compared to the abundant physiological matrix protein that epidermal MCs face (COL IV), COL I and FN ̶ representative of a dermal matrix and thus an altered environment ̶ elicit notable phenotypic shifts in MCs. Relative to COL IV, COL I reduced MC migration and increased melanin production, while FN promoted migration and decreased both proliferation and melanin production. These contrasting effects suggest that ECM composition can selectively modulate distinct aspects of MC behavior. Notably, the intermediate state observed on COL IV supports a model in which this basement membrane component enables MCs to maintain phenotypic flexibility — for example, allowing them to increase melanin production in response to external stimuli such as UV or inflammation. The opposing responses of MCs to FN and COL I underscore the importance of ECM composition in regulating MC function, suggesting that context-dependent ECM remodeling can actively shape MC behavior, with relevance not only for MC homeostasis but potentially also for pathophysiological states and stress responses.

Our findings open the possibility that ECM alterations can disrupt MC homeostasis, with potential beneficial or detrimental consequences for skin health depending on the context. For instance, wound healing reflects a complex physiological process in which MC phenotypic shifts could have a significant impact. Upon skin injury, the initial stage of tissue repair entails the formation of a provisional FN-rich environment (Potekaev et al., 2021), which may facilitate the repopulation of MCs within the regenerated tissue (Snell, 1963). Taking into account our observation that FN enhances MC motility in vitro (fig. 2), it is tempting to speculate that this FN-enriched tissue enables MCs to efficiently migrate into wound sites and re-establish their protective function, such as melanin-based protection from UV-induced damage. Conversely, in fibrotic conditions such as scleroderma, marked by stiffening of the skin due to excessive deposition of COL I, cases of localized hyperpigmentation have been reported in patients, possibly reflecting COL I-driven MC reprogramming. Vitiligo, which is characterized by the loss of epidermal MCs and non-pigmented patches, is associated with a decrease in COL IV and FN and a concomitant increase in COL I content (Rani et al., 2023). The repigmentation process requires MCs to migrate into depigmented areas and synthesize melanin effectively (Norris et al., 1994). Considering our observation of strongly reduced MC migration on COL I, it seems plausible that COL I enrichment in vitiligo lesions interferes with the efficient MC redistribution into depigmented areas in the skin. Furthermore, melasma is a multifactorial skin condition that is characterized by focal hypermelanosis. Sex hormones and UV radiation have been implicated in the development of melasma (Espósito et al., 2022). Interestingly, disruption of the basement membrane has also been reported in 96% of melasma lesions, with MCs protruding into the dermal layer in 66% of cases (Torres-Álvarez et al., 2011). While the mechanism of this dermal invasion by MCs remains open, it has been proposed that the migration of “hyperactive” MCs into the dermis leads to the constant hyperpigmentation in melasma (Phansuk et al., 2022; Torres-Álvarez et al., 2011). This concept would be in line with our findings of COL I-triggered melanogenesis, opening the possibility that MC hyperactivity in melasma could, at least in part, result from the exposure to dermal COL I.

Together, these examples illustrate how context-dependent ECM remodeling could shape MC behavior, with implications not only for physiological repair but also for pathological skin remodeling. Though further investigation is warranted, these observations open new avenues to explore how shifting ECM landscapes influence MC phenotypes in vivo.

### FN-induced phenotypic reprogramming rewires MCs toward a dedifferentiated state

Our study also highlight for the first that ECM components are pivotal in regulating the phenotypic plasticity of MCs. We demonstrate that a FN-rich environment rewires MCs toward a dedifferentiated state, marked by reduced melanin production, slow cycling, and increased motility, while COL I elicits features of a differentiated phenotype, promoting melanin synthesis but limiting migration. In addition to these phenotypic observations, our data indicates that FN exposure induces a distinct transcriptional program associated with MC dedifferentiation. Transcriptomic profiling revealed that FN-cultured MCs downregulate melanocytic differentiation markers and upregulate genes linked to plasticity, stemness, and neural crest-like features. This phenotype is indicative of an adaptive, dedifferentiated state, distinct from the more stable melanogenic profile observed on both COL I and COL IV, and suggests that FN may act as a cue for reprogramming MC identity.

Notably, this dedifferentiated signature observed in FN-exposed MCs is reminiscent of the early phenotypic changes reported during malignant transformation of MCs. Indeed, in a mouse model recapitulating features of human melanomagenesis, it has been shown that mature MCs expressing a B-Raf oncogene can undergo transcriptional reprogramming, associated with a loss of their differentiated characteristics and eventual invasion into the dermis (Köhler et al., 2017). This suggests that MC dedifferentiation precedes MC transformation and melanoma development. Hence, the dedifferentiated phenotype observed in FN-exposed MCs could reflect one of several steps that are required during early stages of melanoma initiation. Taken together, this raises the possibility that a FN-rich environment, combined with oncogenic mutations, could render MCs susceptible to transformation and ultimately melanomagenesis.

In conclusion, our findings underscore the remarkable plasticity of MCs and their ability to shift differentiation states in response to environmental cues, highlighting the importance of ECM molecules in regulating MC behavior. By modulating the balance between melanogenic identity and motile potential, the ECM — particularly FN — may influence not only physiological processes like tissue repair but also pathological trajectories including pigmentation disorders and oncogenesis. Understanding how different conditions elicit these shifts could provide valuable insights into normal skin physiology, the mechanisms underlying melanocyte-related conditions like melanoma, and potential therapeutic targets for restoring normal melanocyte function in pigmentation disorders. In particular, elucidating how ECM remodeling contributes to MC dedifferentiation could open new therapeutic avenues in both regenerative medicine and cancer biology.

### ECM-dependent ERK activation controls MITF localization and activity, and melanogenesis

Given the well-established role of MITF as a master regulator of MC differentiation, pigmentation, and survival, and its known regulation by ERK signaling, we reasoned that it could function as a key integrator of ECM-derived cues. We therefore centered our analysis on MITF, aiming to understand how its localization and activity might be shaped by the extracellular microenvironment. This hypothesis was supported by our finding that ECM-dependent phenotypic shifts of MCs are tightly linked to differential activation of the MEK/ERK pathway, which in turn governs the localization and output of MITF. Specifically, COL I limits ERK activity, triggering high nuclear MITF, while FN stimulates high ERK activity resulting in reduced nuclear MITF levels. This is further associated with adaptation of MC functions under the control of MITF, whereby proliferation and melanin production are enhanced on COL I but reduced on FN. In line with previous studies that linked RAF and MEK/ERK activation to cytoplasmic retention and nuclear export of MITF in melanoma cells, respectively (Estrada et al., 2022; Ngeow et al., 2018), we here demonstrate in MCs that MEK and ERK inhibition reverted the low nuclear MITF and melanin production in FN-exposed MCs. This, together with our findings on ECM-dependent differential ERK activation in MCs, identifies a hitherto unrecognized role of ECM cues in MITF nuclear localization through the regulation of ERK activity. Our data further suggest that FAK may act upstream of ERK, as FAK is more activated on FN and its inhibition on FN reduced ERK activation and melanin production.

Our transcriptomic analysis also pointed to ECM-induced expression changes in Wnt pathway components, e.g. an upregulation of *Frat2* and *Ccnd1* on FN. FRAT2 is known to stabilize β-catenin by inhibiting its GSK3β-mediated degradation (van Amerongen & Berns, 2005), thereby supporting β-catenin–dependent transcription. Since β-catenin can enhance MITF transcription through TCF/LEF-mediated activity (Widlund et al., 2002; Yasumoto et al., 2002), and given that MITF itself can interact with β-catenin to regulate gene expression (Schepsky et al., 2006), these findings open the possibility that Wnt signaling -next to FAK/MEK/ERK signalling-could contribute to ECM-dependent modulation of MITF levels and function.

Overall, further studies are needed to dissect the specific molecular signals downstream of FN that drive the shift of mature MCs toward less differentiated states and to delineate the respective contributions of individual signaling pathways in ECM-mediated responses.

### Interplay between ECM components and substrate stiffness in the mechanosensation of MCs

Our findings underpin a crucial role for ECM components in regulating mechanosensation in MCs and show that ECM components and substrate stiffness govern MC functions. So far, studies have predominantly explored either the roles of different ECM subtypes using ultra-hard substrates like glass or plastic (Hara et al., 1994), or focused on the impact of substrate stiffness with a single ECM type (Choi et al., 2014). In contrast, our approach, combining varying substrate stiffness with different ECM proteins, demonstrated that 1) over the range of stiffness and ECM molecules tested, matrix protein subtypes can dominate over mechanical substrate surface properties, as altering ECM proteins produced more pronounced effects on MC behavior compared to changes in substrate stiffness; and 2) the ability of MCs to respond to mechanical cues is highly dependent on the specific ECM proteins present. Notably, COL I supports stiffness-induced MITF nuclear localization and MC proliferation, while FN enhances mechanosensitive responses at the level of FAK activation and number of FAs. Seong et al. previously reported that in a human fibrosarcoma cell line stiffness-mediated FAK activation can be observed on FN-but not on COL I-coated substrates (Seong et al., 2013). While our observations in MCs are consistent with such effects on FN, iMCs cultured on COL I even showed reduced FAK activity with substrate stiffness, and pMCs had lowest FAK activity at intermediate stiffness. These observations suggest cell type-specific roles of ECM molecules in mechanoresponses.

Strikingly, our data identified MITF as a novel ECM- and mechanosensitive transcription factor, with a stiffness-dependent increase as well as the highest nuclear localization observed on COL I compared to COL IV and FN. Such fine-tuning of MITF subcellular localization may ensure that MC responses are appropriately matched to the environmental matrix protein composition and mechanical features of the ECM. Overall, this highlights the context-dependent nature of MITF regulation downstream of ERK, revealing a hitherto unknown aspect of MITF modulation by ECM cues. Furthermore, our findings suggest that ECM components and substrate stiffness do not merely exert additive effects; instead, individual ECM types demonstrate synergistic interactions with stiffness in regulating MC behavior. This underscores the importance of considering both ECM protein composition and mechanical properties in future studies, as the interplay between these factors significantly influences how cells interpret and respond to mechanical signals, thereby affecting cellular function and differentiation pathways.

## Conclusion

In summary, we report a novel ECM-dependent FAK/MEK/ERK/MITF axis that controls the plasticity of MCs in response to interactions with their environment. Our findings underscore the critical role of the ECM in orchestrating MC function and differentiation, highlighting the need for further studies to fully understand the intricate crosstalk between mechanical parameters and ECM ligands in the cellular microenvironment (fig. 7). Moreover, our data point to a complex link between ECM protein types and substrate stiffness, which can influence MC mechanosensation (fig. 7). Elucidating the specific molecular mechanisms driving this interplay, particularly the roles of MITF and other key regulators, will be crucial for a comprehensive understanding of MC function and pathology.

**Figure 7.**
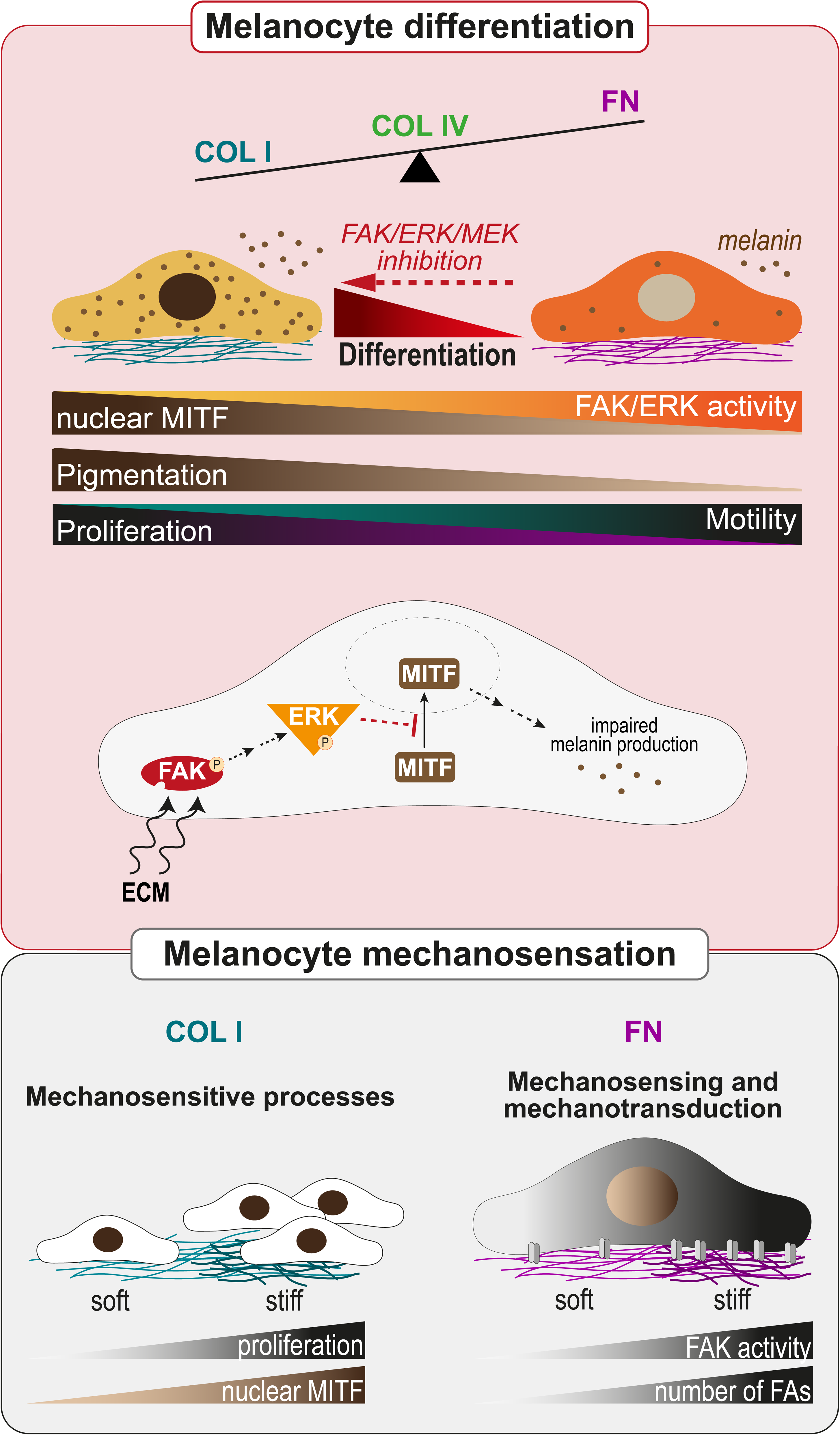
(graphical abstract). Upper panel: MC differentiation is differentially modulated by ECM components. COL I promotes a differentiated phenotype by limiting ERK activation, leading to high nuclear MITF levels, associated with increased pigmentation and proliferation but reduced motility. In contrast, FN rewires MCs towards dedifferentiation, characterized by enhanced ERK activity resulting in reduced nuclear MITF and decreased pigmentation, and associated with a slow-cycling phenotype as well as increased motility. Reducing ERK activity, through FAK, MEK or ERK inhibition, restores the differentiation of MCs exposed to FN. **Lower panel:** ECM components influence how cells interpret and respond to mechanical signals: COL I enables stiffness-mediated MITF nuclear localization and MC proliferation, while FN supports a stiffness-dependent increase in focal adhesion number and FAK activation.

## METHODS

### Preparation of PDMS substrates with tunable stiffness and ECM protein coating

Polydimethylsiloxane (PDMS) substrates were prepared using a SYLGARD 184 Elastomer kit (Dow Silicones, USA) and varying the crosslinker:silicone base agent ratio to 1:60, 1:35, and 1:10, respectively (fig. S1). Once mixed, the PDMS elastomers were cured at 60°C for 16 h. To enhance the hydrophilicity of the substrates and facilitate binding of ECM proteins, all substrates underwent argon plasma treatment using a PDC-002-CE plasma cleaner (Harrick Plasma, USA) at 300 mTorr for 2 min. The substrates were then coated for 2 h at 37°C with 30 µg/ml of mouse collagen IV (Cultrex™, #3410-010-02, Bio-Techne, Germany), 30 µg/ml of type I collagen from calf skin (Collagen G, L7213, Sigma-Aldrich, USA), or 10 µg/ml of bovine plasma fibronectin (F1141, Sigma-Aldrich, USA) diluted in sterile D-PBS (Gibco, USA). After incubation, the excess ECM solution was aspirated, and substrates were washed with sterile D-PBS prior to cell seeding.

### Determination of the elastic modulus of uncoated and coated PDMS surfaces

Atomic Force Microscopy (AFM) served to determine the elastic modulus (Young’s Modulus) of uncoated and protein-coated PDMS surfaces used in this study. For this, the PDMS elastomers (ratio 1:10, 1:35 or 1:60) were prepared in an AFM-suitable imaging dish (ibiTreat µ-Dish, Ibidi, Germany) and transferred to the sample stage of the microscope (Nanowizard 4, Bruker Nano GmbH, Germany). An AFM cantilever with a spherical tip (Biosphere B2000, Nanotools, Germany) with a nominal spring constant of 0.2 N/m and a sphere radius of 2 µm was used to perform nanoindentation measurements. AFM force-distance measurements were conducted in a grid of 4 x 4 points with a lateral distance of 4 µm (= one tip diameter) between two points to probe native spots. The AFM cantilever was extended to the PDMS surface with a speed of 1 µm/s and pressed on it with a loading force of 20 nN. Afterwards, the cantilever was retracted with 1 µm/s to complete the force-distance measurements. For each force-distance curve, the Hertz fit (Horvath et al., 2019) was applied to the contact region of the extend curve to evaluate the Young’s Modulus with the JPK SPM Data Processing Software v8.0.1 (Bruker Nano GmbH, Germany).

### Mice

All mice were housed and maintained according to federal guidelines within the specific pathogen free (SPF) animal facility in Building 61.4 of the Saarland University in Homburg under the breeding license of Sandra Iden’s lab (§11 Az. 2.4.1.1-Iden). All experiments were performed in line with German animal welfare laws as well as institutional guidelines and were reported to the responsible authorities (Vorabmeldung VM2023-16 and previous notifications). Wildtype (wt) mice were either purchased from the Jackson Laboratory (Charles River Germany) or were taken from in-house breedings of transgenic mouse lines. In those cases,

F1 and F2 generation were genotyped to verify the wt alleles of the genes of interest of the respective mouse line.

### Isolation and culture of primary mouse melanocytes

Primary mouse MCs were isolated from the epidermis of newborn wt mice and maintained up to passage 3 as previously described (Mescher et al., 2017). Briefly, to separate epidermis from dermis, whole skin from mice at postnatal day (P)0-P3 were incubated in a solution of 5 mg/ml Dispase II (#D4693, Sigma-Aldrich, USA) diluted in RPMI medium (Gibco, USA) supplemented with 10% fetal calf serum (FCS, #S0615, Sigma-Aldrich, USA), penicillin (100 U/ml), streptomycin (100 μg/ml), 100 nM sodium pyruvate, and 10 mM non-essential amino acids (all Gibco, USA) at 4°C overnight. The next day, the epidermis was incubated for 20 min in TrypLE select (#12563-011, Gibco, USA) at room temperature. Dissociated cells were collected and cultured in RPMI medium containing 200 nM TPA (12-O-tetradecanoylphorbol-13-acetate, #P8139, Sigma-Aldrich, USA) and 200 pM cholera toxin (#C8052, Sigma-Aldrich, USA), hereafter referred to as RPMI+. At passage 0, the cell cultures contained non pigmented melanoblasts, MCs, and keratinocytes. After 7 days, cells were passaged, and due to terminal differentiation, keratinocytes were largely removed, resulting in MC monocultures suitable for further experiments.

### Generation of stable murine melanocyte cell lines (iMCs)

Immortalized MCs (iMCs) were generated by immortalizing mouse primary MCs (mTmG-positive) with SV40 large T antigen, followed by subcloning. The cells express membrane-targeted tandem dimer Tomato (dt; Muzumdar et al., 2007), yielding red fluorescence. iMCs exhibit morphological and molecular characteristics common to normal murine skin MCs. iMCs were maintained in culture in RPMI+ at 37°C with 5% CO_2_.

### Immunofluorescence

To assess cell area and morphology, and for quantification of p-FAK, Ki67, MITF, YAP and p-ERK signal intensities, MCs were seeded on the different substrate combinations at a density of 2,500 cells/cm², to avoid overcrowding and allow single cell measurements. 24 h later, MCs were fixed in 4% PFA/PBS for 10 min. For MITF, YAP and p-FAK stainings, cells were blocked for 1 h at room temperature with 5% bovine serum albumin (BSA) in TBS containing 0.2% Triton X-100 (named blocking buffer hereafter). Primary antibody incubation was performed in blocking buffer diluted at a ratio of 1:5 in TBS at 4°C overnight. For Ki67 and p-ERK stainings, blocking and primary antibody dilution were performed using PB buffer composed of 0.05% milk powder (#T145.2, Carl Roth, Germany), 0.25% fish gelatin (#G7765, Sigma-Aldrich, USA), 0.5% Triton X-100 (#BP151-100, Fisher Bioreagents, Germany), 20 mM HEPES (pH 7.2, #9105.4, Carl Roth, Germany), and 0.9% NaCl (#3957.1, Carl Roth, Germany). For all, after 3x washing in 0.2% Tween/TBS, MCs were stained with AlexaFluor-conjugated secondary antibodies, DAPI (Carl Roth, Germany) and phalloidin to stain actin for pMCs for 1 h at room temperature. After washing, PDMS scaffolds were mounted onto glass coverslips using Fluoromount-G™ (#00-4958-02, ThermoFisher Scientific, USA). Antibodies used for immunostainings are listed in the Antibodies and reagents section.

### BrdU Assay

48 h post-plating, MCs were incubated with 160 µg/ml of BrdU (5-Brom-2′-desoxyuridin; #10280879001, Roche, Germany) for 2 h at 37°C and then washed with PBS and fixed with 4% PFA for 10 min at room temperature. Cells were treated with 2 N HCl for 10 min at room temperature to achieve DNA denaturation, followed by a PBS wash and subsequent immunofluorescence staining as previously described. BrdU-positive MCs were visualized using fluorescence microscopy (see below) and quantified from at least 20 randomly positioned micrographs. The background signal was subtracted from the measured signal of each nucleus to identify BrdU-positive cells. The proliferation rate was then determined by counting BrdU-positive cells and dividing this number by the total number of cells, identified via DAPI staining.

### Microscopy

Epifluorescence micrographs were acquired with an AxioObserver Z1 inverted microscope equipped with a Colibri 7 LED light source (Zeiss, Germany) using a Plan-Apochromat 20x/0.8 air objective an EC Plan Neofluar 40x/1.3 oil objective, a Plan-Apochromat 63x/1.4 oil objective and an Axiocam 305 camera (Zeiss, Germany) or Rolera ECM2 camera coupled to the ZEN Blue imaging software V2.6 (Zeiss, Germany).

### Image quantification

Image analysis was performed using Fiji/ImageJ software (Schindelin et al., 2012; Schneider et al., 2012) to quantify cell area, morphology, FAK phosphorylation and focal adhesion numbers, ERK phosphorylation, as well as YAP and MITF nuclear translocation in MCs 24 h post-plating.

#### MC dendricity and cell area

Epifluorescence micrographs of actin (phalloidin staining) or td-Tomato, for pMCs and iMCs respectively, were preprocessed using “mean” filter to smoothen images, by reducing noise and averaging pixel values, followed by the “median” filter, which further reduced noise while preserving edges and details. Images were then converted into binary images by using the “Threshold” function to obtain a mask of the cell shape. The binary mask was used either for cell area quantification or for the quantitative assessment of morphological characteristics of MCs. Using the “Sholl Analysis” plugin, concentric circles (named radius), were drawn on the binary image at regular intervals (5 µm) from the center of the cell body. The number of intersections between dendrites and each radius was counted and then plotted to create a Sholl profile, which shows the number of dendrite intersections at increasing distances from the cell body (Binley et al., 2014; Sholl, 1953).

#### p-FAK immunoreactivity

To quantify p-FAK immunoreactivity in MCs, actin (phalloidin staining) or td-Tomato signals were used to detect the cell contour of pMCs and iMCs, respectively. This contour was then superimposed on p-FAK signal, followed by measurement of the integrated density of the signal of individual MCs. For iMCs, data were pooled after normalization for each experiment as two different microscope cameras have been used for these experimental series.

#### Focal adhesion number

To quantify the number of focal adhesions based on p-FAK signals, first, a background subtraction was performed using a rolling ball algorithm. Then, a suitable threshold was applied to isolate the focal adhesion signal from the background. Individual focal adhesions were detected using the “analyze particles” function (0.05-5 µm). Finally, the number of focal adhesions per cell and per unit area was determined.

#### YAP and MITF subcellullar localization

Immunodetection of MITF and YAP were performed in separate experiments and analyzed independently. For both stainings, appropriate negative controls were included (secondary antibody alone without primary antibody), which showed no detectable signal.

To evaluate YAP nuclear localization in response to ECM cues, DAPI was used to delineate nuclear contours, which were overlaid onto the YAP signal to quantify nuclear integrated intensity in individual MC. To account for signal variation across experiments and due to the use of two different cameras along this experimental series, values were normalized within each independent experiment by dividing each individual measurement by the mean nuclear YAP intensity across all conditions.

For MITF quantification, phalloidin-based F-actin staining (in case of pMCs) or td-Tomato signal (for iMCs) was employed to delineate whole-cell contours and DAPI served as nuclear counterstain. These masks were overlaid onto the MITF signal to extract integrated intensity values from the nuclear and whole-cell compartments of individual MCs. Cytoplasmic MITF intensity was calculated by subtracting the nuclear signal from the whole-cell signal. Then, nuclear and cytoplasmic intensity values were normalized within each independent experiment by dividing the values of each individual measurement by the mean total intensity across all conditions, thereby normalizing for inter-experimental variability while preserving relative differences between ECM conditions. Finally, MITF nuclear localization was expressed as the percentage of nuclear signal relative to total cellular signal (value for immunoreactivity in the nucleus/ value for immunoreactivity in the whole cell × 100), allowing comparison of nuclear enrichment across ECM conditions.

#### p-ERK immunoreactivity

To quantify p-ERK immunoreactivity in iMCs, the td-Tomato signal was used to detect the cell contour, which was then superimposed on the p-ERK immunostaining signal, followed by measurement of the integrated density of the signal of individual iMCs. Values were normalized within each independent experiment by dividing the values of each individual measurement by the mean nuclear intensity across all conditions. This approach allowed us to account for variability between experiments while preserving relative differences between ECM conditions.

### Live-cell imaging for analysis of MC motility

Live-cell imaging was conducted to investigate dynamic cellular behaviors in response to ECM. Taking advantage of the stable expression of the fluorescent membrane-targeted Tomato, MCs were seeded at a density of 1,000 cells/cm² in plastic chamber slide coated with the distinct ECM proteins (µ-Slide 8 Well, Ibidi, Germany), and placed in a live cell imaging system (Zeiss CellDiscoverer 7) equipped with a temperature- and CO_2_-controlled chamber. Time-lapse images were acquired every 20 min for 20 h using the epifluorescence microscopy mode. Two methodologies were employed to enable real-time dynamic visualization and quantitative analysis of cell migration. To quantify migration parameters, i.e., total distance migrated by cells, and their speed, cells were tracked using the “Manual tracking” plugin in Fiji software. Additionally, the “Temporal Color Code” in Fiji software was applied to track cell shape and movement over time, allowing to visualize changes of MC dynamics over time. Briefly, different colors represent the shape and position of cells at each time point, with the color gradient indicating the progression of time. The overlay of different colors provides information on the dynamic behavior of cells throughout the observation period.

### RT-qPCR

For quantitative RT-PCRs, RNA was isolated from MCs using TRIzol reagent according to the manufacturer’s protocol (Invitrogen#15596026, ThermoFisher Scientific, USA). RNA was reversely transcribed using a QuantiTect Reverse Transcription kit (Qiagen, Germany) and amplified with iTaq Universal Probes Supermix (Biorad, USA). Target gene expression was detected using TaqMan probes (GAPDH mouse Mm99999915_g1; Tyr mouse Mm00495817_m1; ThermoFisher Scientific, USA). Gene expression changes were calculated using the comparative CT (ΔΔCT) method, normalized to GAPDH expression, and compared to cell lysates from soft substrates coated with COL IV.

### Melanin content assay

The melanin assay was performed to quantify the average melanin amount produced by a cell. Both intracellular melanin and extracellular melanin content (i.e., melanin released into the medium) were measured from the same batch of cells. For melanin assays, each condition was performed in duplicate: one well was used for melanin extraction, and the other for cell counting. MCs were cultured in 12-well plates for 72 h in 1 mL phenol red-free RPMI medium (Gibco, USA) to avoid interference with extracellular melanin measurements. The medium was then collected and analyzed to measure extracellular melanin content. Intracellular melanin was quantified following an adapted protocol from Ito and Wakamatsu (Ito & Wakamatsu, 2003). MCs were lysed in 400 µL of 10% DMSO in 1 N NaOH (extraction buffer) and incubated with agitation (24 h at 100°C for pMCs and 18 h at 80°C for iMCs). Extracellular extracts were quickly centrifuged for 10 sec to sediment cellular debris, and intracellular extracts were centrifuged for 5 min at 13.000 rpm. Extra- and intracellular extracts and their respective blanks, RPMI media or lysis buffer, were loaded onto a 96-well plate and absorbance was measured at 405 nm using a Multiskan™ FC Microplate Photometer (ThermoFisher Scientific, USA). For analysis, the OD values were subtracted by the blank to remove background signals caused by the medium or lysis buffer. Next, extracellular values were multiplied by 5 and intracellular by 2 to account for the stock volume, and the values were divided by the cell number to obtain the relative melanin content per cell.

For intracellular melanin quantification following MITF depletion, iMCs were transfected with 50 nM siMITF (ON-TARGETplus Mouse Mitf siRNA SMARTPool, L-047441-00-0005, Dharmacon, UK) or control siRNA (siGENOME Control Pool, D-001206-14-20, Dharmacon, UK) using the Viromer® Blue kit (Lipocalyx) 4 h after plating. Cells were harvested and lysed for melanin extraction 72 h post-transfection using the same procedure as above.

### Transcriptomic analysis

#### RNA extraction, quality control, library preparation, and sequencing

Total RNA was extracted using TRIzol reagent (Invitrogen, ThermoFisher Scientific, USA) or the RNeasy Mini Kit (Qiagen, Germany), following the manufacturer’s instructions. For each condition, RNA was prepared from three biological replicates. RNA quantity and purity were assessed using a NanoDrop One© spectrophotometer (ThermoFisher Scientific, USA), and RNA integrity was verified using the RNA Integrity Number (RIN). Only samples with RIN values ≥4 were processed further. As additional quality controls, melanin content assays and RT-qPCR were performed in parallel, targeting melanocyte-specific markers *Mitf* and *Tyr*.

RNA sequencing was performed by Novogene (Munich, Germany) using standard Illumina protocols. Polyadenylated mRNAs were enriched, fragmented, and reverse-transcribed into cDNA. Strand-specific libraries were prepared and quality-checked using Qubit, real-time PCR, and a Bioanalyzer, then sequenced on Illumina platforms (2 × 150 bp), generating approximately 25 million paired-end reads per sample. Raw reads were processed with fastp to remove adapters and low-quality reads. Clean reads were aligned to the mouse genome (GRCm39/mm39) using HISAT2, and gene expression was quantified with featureCounts and normalized as FPKM values. RNAseq data have been deposited at GEO (accession GSE297747).

#### Bioinformatics analysis

Gene expression data from cells cultured on COL1, COL4, and FN were processed using Pandas (McKinney, 2010) and NumPy (Harris et al., 2020) for sorting and initial exploration. Differential gene expression analysis was performed using the Bioconductor package DESeq2 (Love et al., 2014) to identify genes significantly regulated between the conditions. Genes with an adjusted p-value < 0.05 and |log2 fold change| ≥ 1 were considered differentially expressed. Gene expression values were Euclidean-normalized, and heatmaps were created using Seaborn (Waskom, 2021) to visualize expression patterns across conditions. Results were visualized using volcano plots generated with Matplotlib (Hunter, 2007). To explore biological relevance, association of significantly regulated genes with biochemical pathways was analyzed using Reactome Pathway (Milacic et al., 2024), KEGG Pathway (Kanehisa et al., 2025), and WikiPathways (Agrawal et al., 2024). In addition, Gene Ontology (GO) (Thomas et al., 2022) enrichment analysis was performed to identify overrepresented biological processes, providing insight into the functional impact of ECM proteins on gene regulation.

### Preparation of cell extracts, SDS-PAGE, and immunoblotting

MCs cultures were lysed using boiled SDS/EDTA solution (1% SDS, 10 mM EDTA) and genomic DNA was sheared by passing the lysates through a 27 G x ¾ canula. Protein concentrations were quantified using the BCA assay according to the manufacturer’s protocol (Pierce, ThermoFisher Scientific, USA). SDS-PAGE was carried out using 8-10% polyacrylamide gels. After completing gel electrophoresis, the proteins were transferred onto a PVDF membrane at 20 V for 60 min. Following the transfer, the membrane was blocked for 1 h at room temperature using 5% BSA/TBST (TBS containing 0.2% Tween 20). The primary antibody was incubated in 5% BSA/TBST overnight at 4°C on a roller. Membranes were then washed 3x for 10 min with TBST and incubated with an HRP-conjugated secondary antibody in 5% BSA/TBST for 1 h at room temperature on a roller. Membranes were washed 3x for 10 min with TBST, excess TBST was removed, and membranes were incubated 1 min with in a 1:1 mixture of ECL solutions A and B (PerkinElmer, USA). Excess ECL solution was removed, and the membrane was imaged using the Amersham ImageQuant^TM^ 800 detector (Cytiva, USA). Following a 30-min wash at room temperature with a buffer containing 0.2 M glycine and 0.1% SDS (pH 2.5) to strip the membranes, they were blocked and immunoblotted for total ERK using the same membranes previously used for phospho-ERK detection.

### Quantification of phospho-ERK signal in immunoblot analyses

The protein band intensity of non-saturated Western blot signals was quantified using Fiji software. Each band was selected on the original .tiff image obtained with the Amersham ImageQuant^TM^ 800 Detector, and the “Plot Lanes” tool generated intensity profiles. Peaks corresponding to individual bands (related to total or phosphorylated protein) were identified, and the area under them was measured. The p-ERK/total ERK ratio was calculated to assess relative ERK activity under the different conditions. The values were normalized to the sum of each membrane’s values to determine the mean and standard deviation across experiments.

### Inhibitor treatment

To assess the role of ERK activity for ECM-dependent MC functions, MCs were seeded 4 h prior a 24h- or 72h-treatment with 100 nM Trametinib (MEK inhibitor, #10999, Biozol, Germany), 10 µM of Ravoxertinib (ERK inhibitor, #S7554, Selleckchem, Germany) or 10 µM of Ifebemtinib (FAK inhibitor, #E1114, Selleckchem, Germany) diluted in RPMI+.

### Antibodies

Details on antibodies and dyes used in this study are listed in the following Table S1.

### Statistical analyses

Statistical analysis was performed using GraphPad Prism software (GraphPad, version 10.2.1). Measures of pooled data, from at least three independent experiments, are represented by mean and SD or SEM, as indicated in the figure legends. All data sets were subjected to normality tests (Anderson-Darling test, D’Agostino & Pearson test, Kolmogorov-Smirnov test, or Shapiro-Wilk test) when applicable. Then, statistical significance was determined using One-way ANOVA with Welch’s correction (unequal variances) with Dunnett’s T3 post-hoc test; RM one-way ANOVA with Geisser-Greenhouse’s correction (unequal variances) with Tukey’s post-hoc test; Ordinary two-way ANOVA Tukey’s post-hoc test; Multiple t tests with the Holm-Sidak method’s correction with Dunn’s post-hoc test; and for non-Gaussian datasets using Kruskal-Wallis test with Dunn’s post-hoc test, as indicated in the figure legends. P-values are ranged as follows: *, P < 0.05; **, P < 0.01; ***, P < 0.001; and ****, P < 0.0001, as detailed in the Table S2. The number of independent biological replicates and number of single cells analyzed is specified in each figure legend and in the Table S2.

### Software

Data collection used the following software: Microscopy: ZEN Blue 2.6 (Zeiss, Germany); Immunoblot: Amersham ImageQuant^TM^ 800 control software (Cytiva, USA); qRT-PCR: CFX Opus 96 (BioRad, USA). For data analysis, the following softwares were used: GraphPad PRISM, version 10.2.1, ImageJ/Fiji, JPK SPM Data Processing Software v8.0.1 (Bruker, Nano, Germany).

## ACKNOWLEDGEMENTS

We thank the animal facility of the Faculty of Medicine, Saarland University, for important services, the Robert Ernst lab (Medical Biochemistry and Molecular Biology) for access to the AxioObserver Z1 microscope, the Department of Cellular Neurophysiology (CIPMM, Saarland University) for kindly providing access to the plasma cleaner, and Dr. Nicole Ludwig for access to a TapeStation for RIN measurements. We further thank all members of the Iden laboratory for stimulating discussions.

## DATA AVAILABILITY

Correspondence and requests for materials related to this study should be sent to. All data supporting the findings of this study are available within the paper and its supplemental information, or from the corresponding author on reasonable request.

## AUTHOR CONTRIBUTIONS

**Conceptualization and Supervision**: C.L., S.I.; **Methodology**: C.L., M.D., E.B., V.S.R., G.G., S.T., A.K.B., A.R., S.I.; **Validation**: C.L., M.D., E.B., V.S.R., G.G., V.H., S.I.; **Investigation**: C.L., M.D., V.S.R., E.B., N.D., E.W., G.G., S.T.; **Formal analysis**: C.L., M.D., V.S.R., E.B., G.G., S.T.; **Visualization**: C.L., S.T., S.I.; **Resources**: A.K.B., A.R., V.H., M.B., S.I.; **Writing - first concept:** C.L.; **Writing - original draft**: C.L., S.I.; **Review and editing of manuscript**: C.L., S.I., A.K.B., V.R:, M.B, V.H.; and **Funding acquisition**: S.I.

---

**Figure S1:**
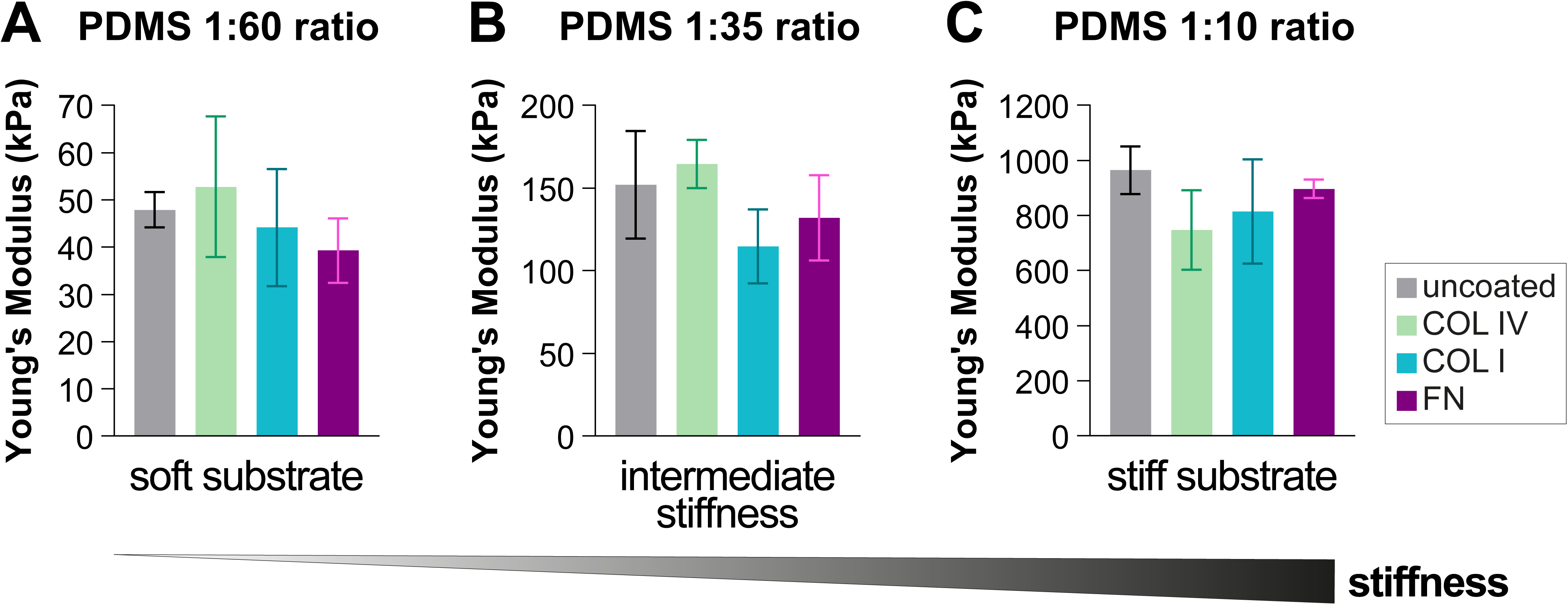
Validation of PDMS elasticity using Atomic Force Microscopy. Young’s modulus of PDMS substrates with varying crosslinker ratios. **A**. PDMS substrate with a crosslinker:silicone base agent ratio of 1:60; means±SD, N=3; Welch ANOVA tests: ns, p>0.05. **B**. PDMS substrate with a 1:35 ratio; means±SD, N≥3; Welch ANOVA tests: n.s.: p>0.05. **C**. PDMS substrate with a 1:10 ratio; means±SD, N=3; Welch ANOVA tests: n.s.: p>0.05.

**Figure S2:**
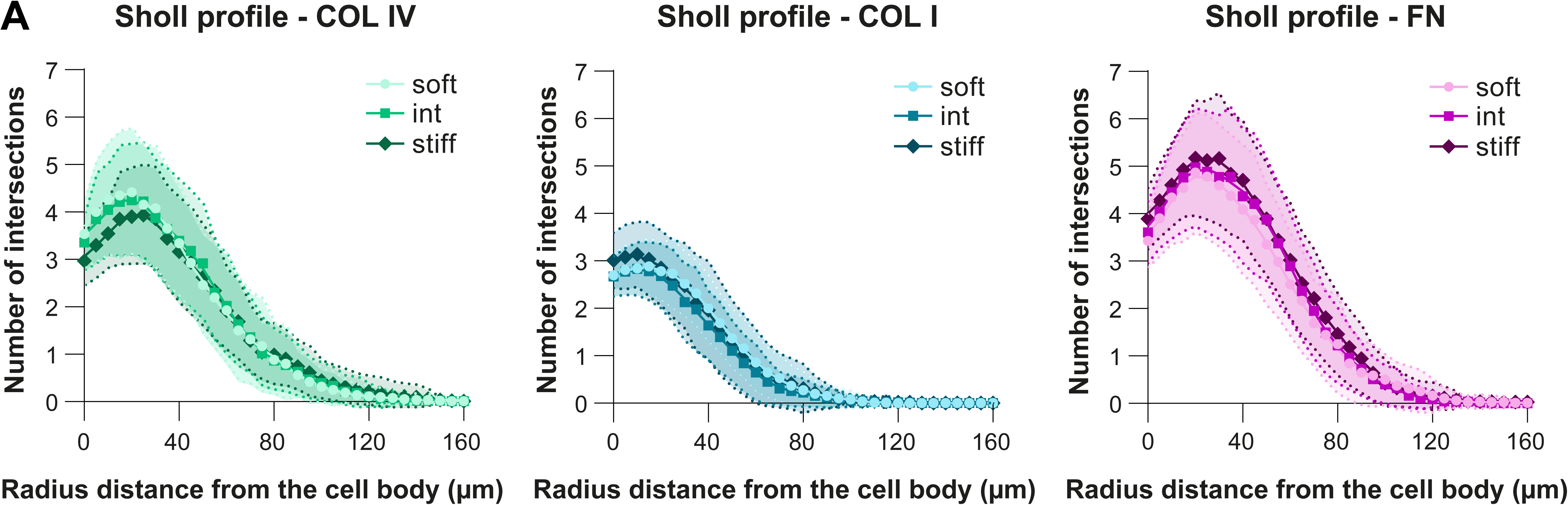
Sholl profiles for iMCs exposed to different ECM molecules. **A.** Graphs showing the Sholl profile analysis, plotting the number of dendrite intersections against the distance from the cell body. The SEM are represented by the connecting curve (dotted line); N=3, n(cells)≥85. Each curve represents a substrate stiffness condition, and each graph represents an ECM type (same cells as analyzed in fig. 1C, different plotting of the data); Welch ANOVA test: n.s.: p>0.05.

**Figure S3:**
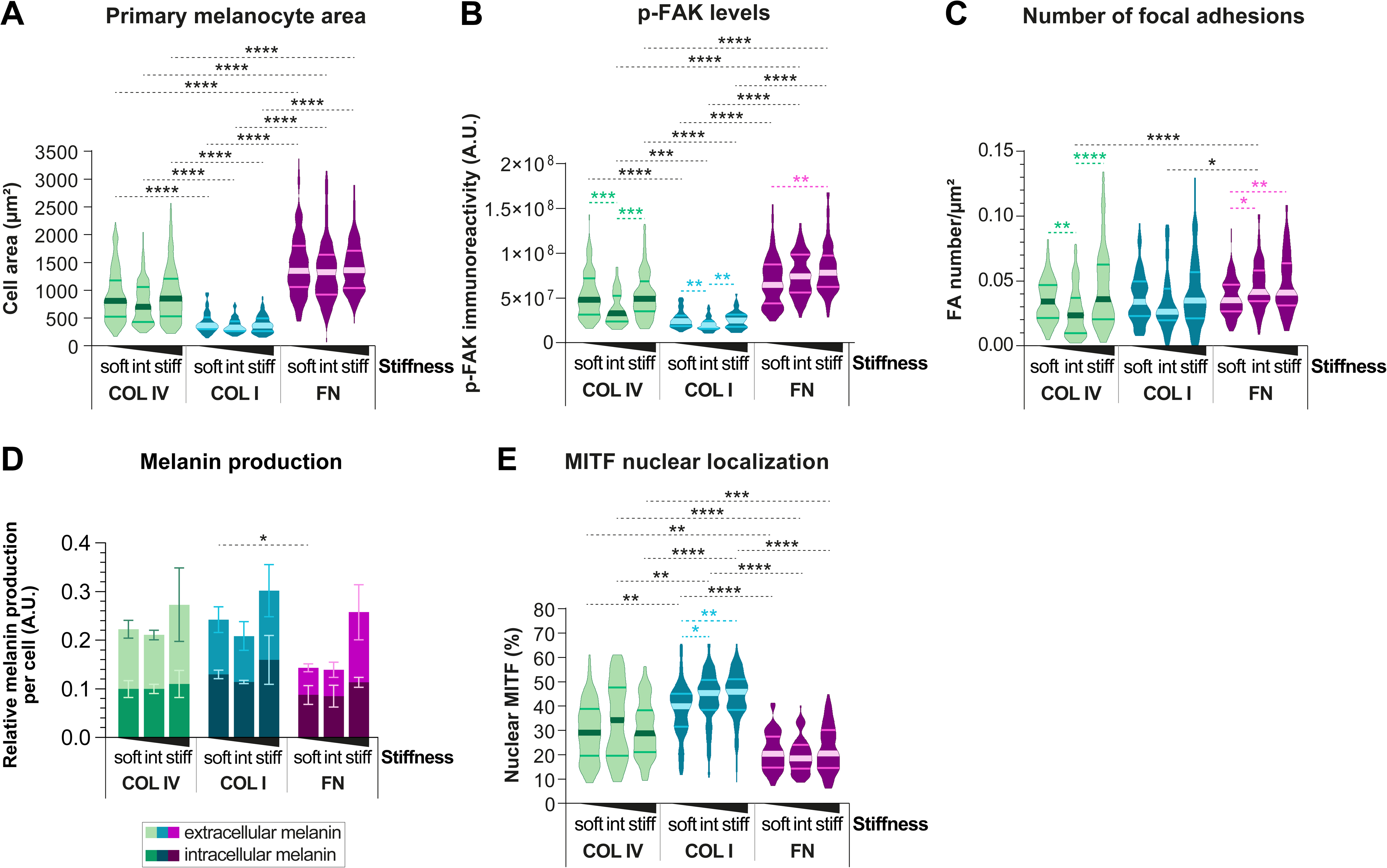
Response of primary MCs to distinct ECM cues. pMCs were cultured overnight on substrates of varying stiffness (soft, intermediate or stiff) coated with COL IV, COL I or FN. **A.** Violin plots showing the medians and distributions of the cell area of pMCs; N≥5, n(cells)≥96; Kruskal–Wallis test: ****: p<0.0001. **B.** Quantification of p-FAK levels; violin plots showing the medians and distributions of the integrated density of total p-FAK per cell; N=3, n(cells)≥52; Kruskal–Wallis test: ****: p<0.0001; ***: p<0.001; **: p<0.005. **C.** Quantification of FAs; violin plots showing the medians and distributions of the number of focal adhesions per µm^2^ per cell; N=3, n(cells)≥65; Kruskal–Wallis test: ****: p<0.0001; **: p<0.005; *: p<0.05. **D.** Quantification of intra- and extracellular melanin content by spectrophotometry at 405 nm from pMCs cultured 72 h on substrates of varying stiffness (soft, intermediate or stiff) coated with COL IV, COL I or FN; means±SD, N=3; multiple t tests: *: p<0.05. **E.** Quantification of nuclear MITF; violin plots showing the medians and distributions of the percentage of nuclear MITF per cell; N=3, n(cells)≥67; Kruskal–Wallis test: ****: p<0.0001; ***: p<0.001; **: p<0.005; *: p<0.05. Abbreviations: A.U., arbitrary units.

**Figure S4:**
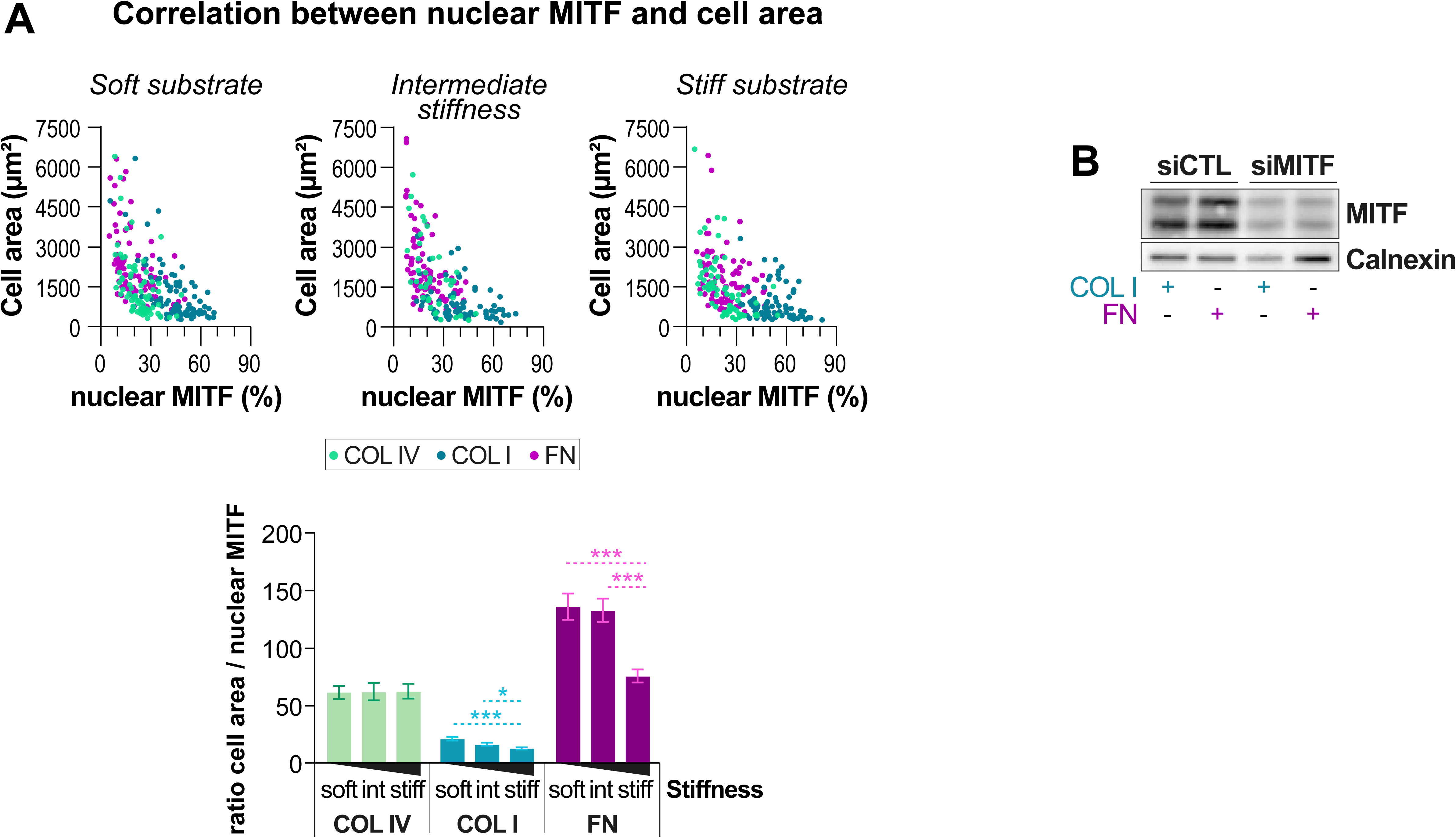
Negative correlation between nuclear MITF levels and MC area and siMITF validation. **A**. The percentage of nuclear MITF was plotted against the cell area of the corresponding MC for all stiffnesses and ECM types tested. The bar diagram depicts the means±SEM of the ratio of cell area to percentage of nuclear MITF per cell; N=3, n(cells)≥47; Kruskal–Wallis test: ****: p<0.0001; ***: p<0.001; **: p<0.005; *: p<0.05. **B.** Western blot analysis of MITF expression in iMCs plated on COL I or FN and transfected for 72 hours with siCtrl or siMITF, performed in parallel with melanin quantification shown in Fig. 4D.

**Figure S5:**
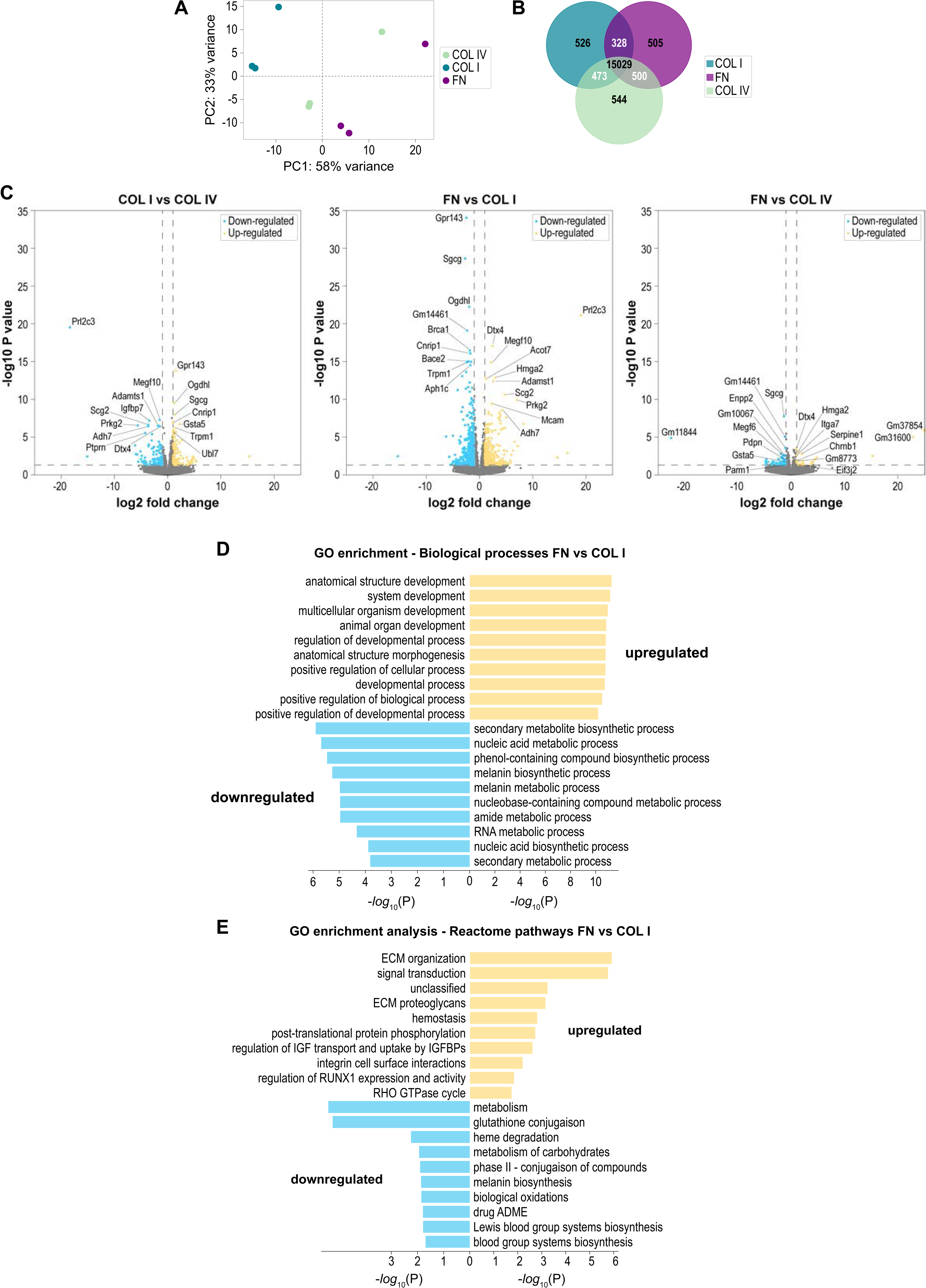
Transcriptomic profiling across ECM conditions. **A.** Principal Component Analysis (PCA) of gene expression across substrates. Each dot represents one sample: COL IV (green), COL I (blue), and FN (purple). Axes indicate principal components capturing the highest variance. **B.** Venn diagram showing the number of genes expressed in the samples grown on COL IV, COL I and FN substrates. **C.** Volcano plots depicting differentially expressed genes between substrates: (i) COL I vs. COL IV, (ii) FN vs. COL I, and (iii) FN vs. COL IV. Significantly downregulated genes are shown in blue; upregulated genes in yellow. **D.** Gene Ontology (GO) Biological Process enrichment analysis of significantly regulated genes across substrates. The x-axis indicates the adjusted p-value of Fisher’s Exact test; the y-axis lists the most significantly enriched biological processes. **E.** Reactome pathway enrichment analysis performed on differentially expressed genes across ECM conditions. The x-axis represents the adjusted p-value of Fisher’s Exact test; the y-axis lists the top significantly enriched Reactome pathways.

**Figure S6:**
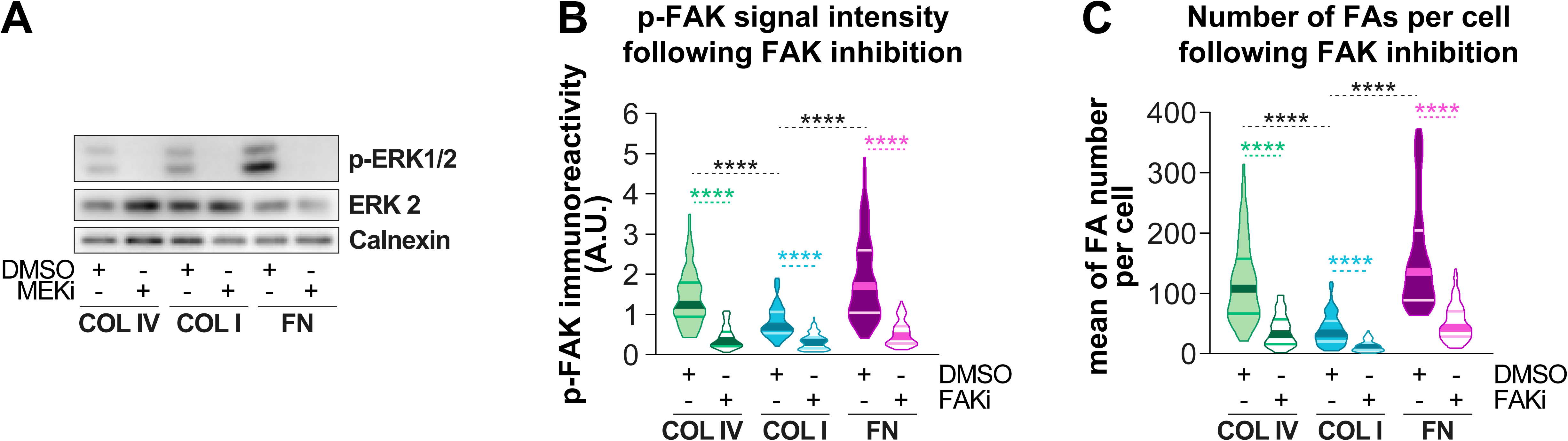
Functional validation of MEK and FAK inhibitors in iMCs cultured on ECM-coated substrates. iMCs were cultured on stiff substrates coated with COL IV, COL I or FN. 4 h post-plating, a 16-hrs treatment with DMSO, 100 nM of MEK inhibitor (MEKi = Trametinib) or FAK inhibitor (FAKi = Ifebemtinib) was commenced. **A.** Representative Western blot showing the inhibition of ERK1/2 phosphorylation with Trametinib treatment. **B**. Quantification of p-FAK (Y397) intensity per cell following treatment with DMSO or FAK inhibitor (FAKi); violin plots display median and distribution; N=3, n(cells)≥88; Kruskal–Wallis test: ****: p<0.0001. **C**. Quantification of focal adhesion (FA) number per cell following FAKi treatment; violin plots show median and distribution; N=3, n(cells)≥70; Kruskal–Wallis test: ****: p<0.0001.

## Supplementary tables

**Supplementary Table S1.**

| **Primary Antibodies** |  | **Company** | **Catalog Nr.** |
| --- | --- | --- | --- |
| **IF** | Anti-MITF antibody [C5] | Abcam | Ab12039 |
|  | Phospho-FAK (Tyr397) antibody | Cell Signaling Technology | 3283S |
|  | YAP (D8H1X) XP® | Cell Signaling Technology | 14074S |
|  | BrdU | Biolegend | 339802 |
|  | Ki67 | Biolegend | 652401 |
|  | pERK1/2 | Cell Signaling Technology | 4370 |
| **WB** | pERK1/2 | Cell Signaling Technology | 4370 |
|  | ERK2 | BD Pharmingen | 610104 |
|  | MITF | Cell Signaling | 97800 |
|  | Calnexin | Enzo Life Sciences | ADI-SPA-860-D |
| **Conjugated Secondary Antibodies** |  |  |  |
| **IF** | Alexa Fluor™ 488 donkey anti-rabbit | Invitrogen | A-21206 |
|  | Alexa Fluor™ 647 goat anti-mouse | Invitrogen | A-21235 |
| **WB** | HRP donkey anti-rabbit | Invitrogen | A16023 |
|  | HRP sheep anti-mouse | Amersham | NA931V |
| **Dyes** |  |  |  |
| **IF** | Alexa Fluor™ 568 Phalloidin | Molecular Probes | A12380 |
|  | DAPI | Carl Roth | 6335.1 |

**Supplementary Table S2.**
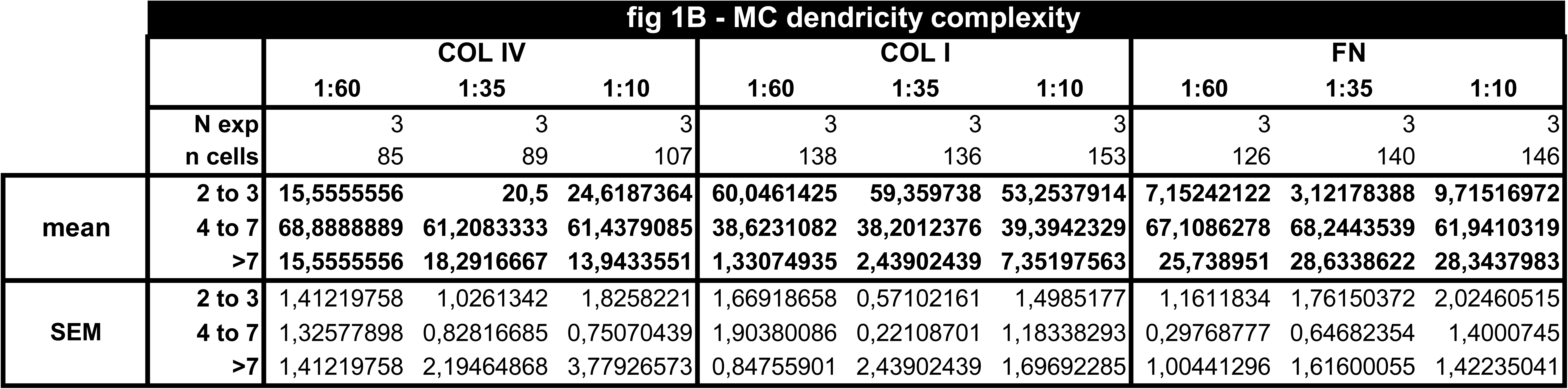
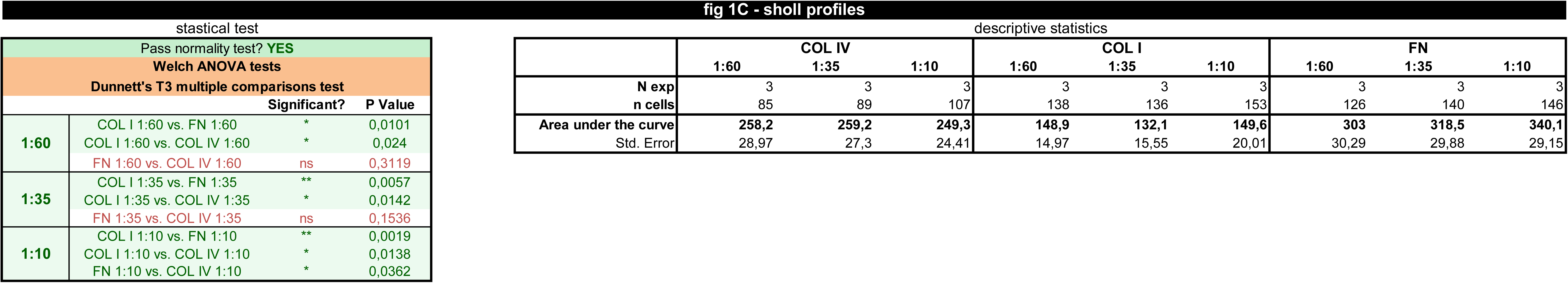
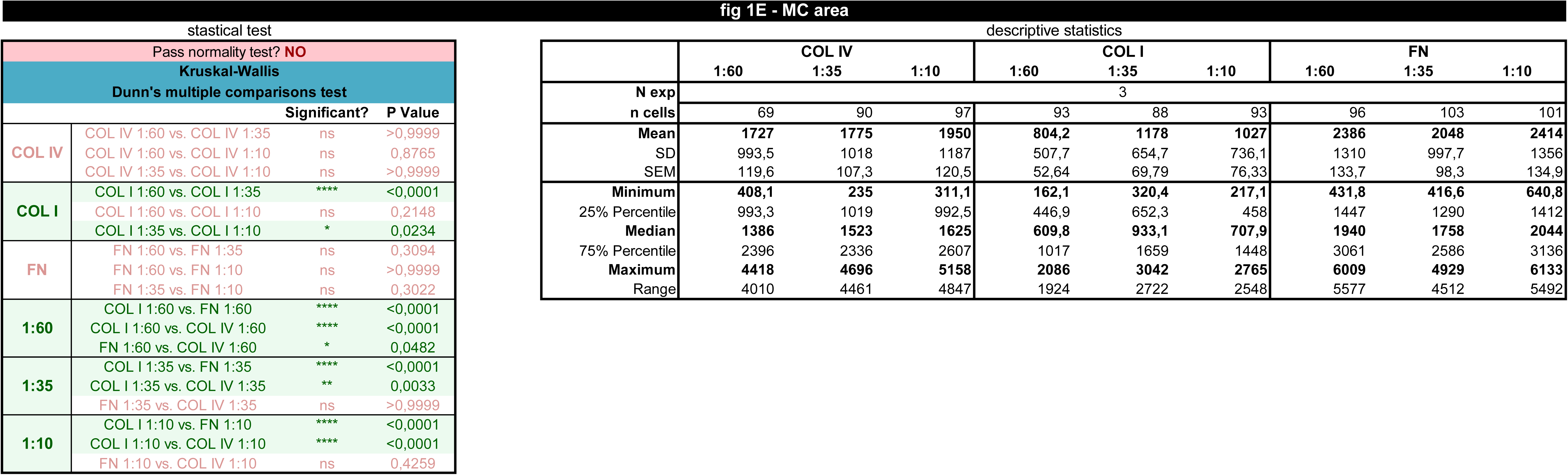
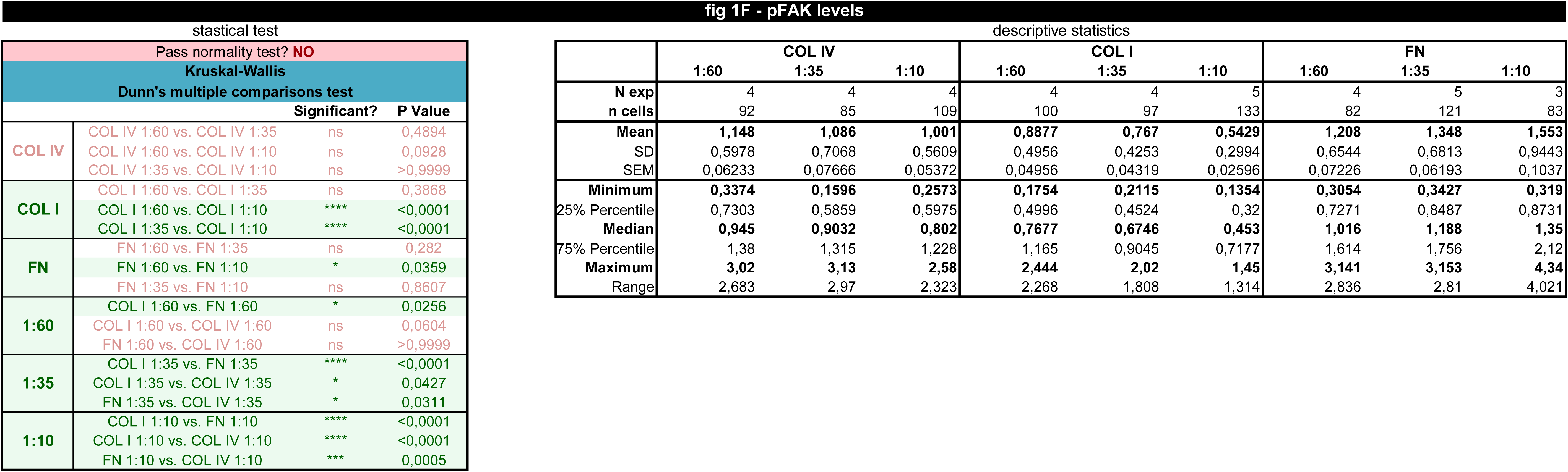
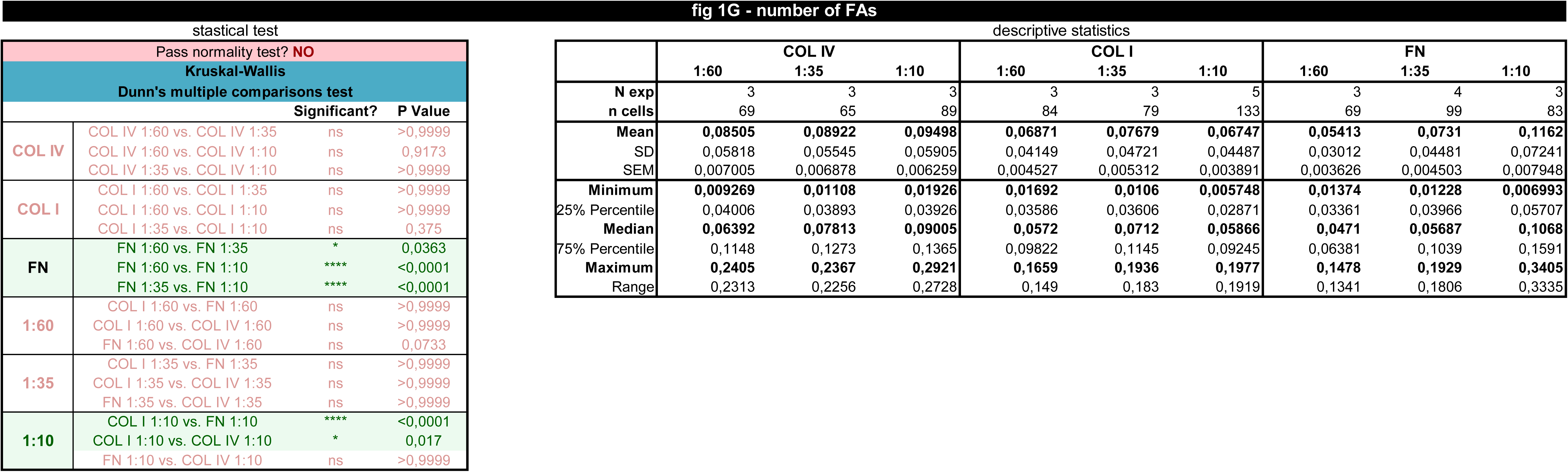
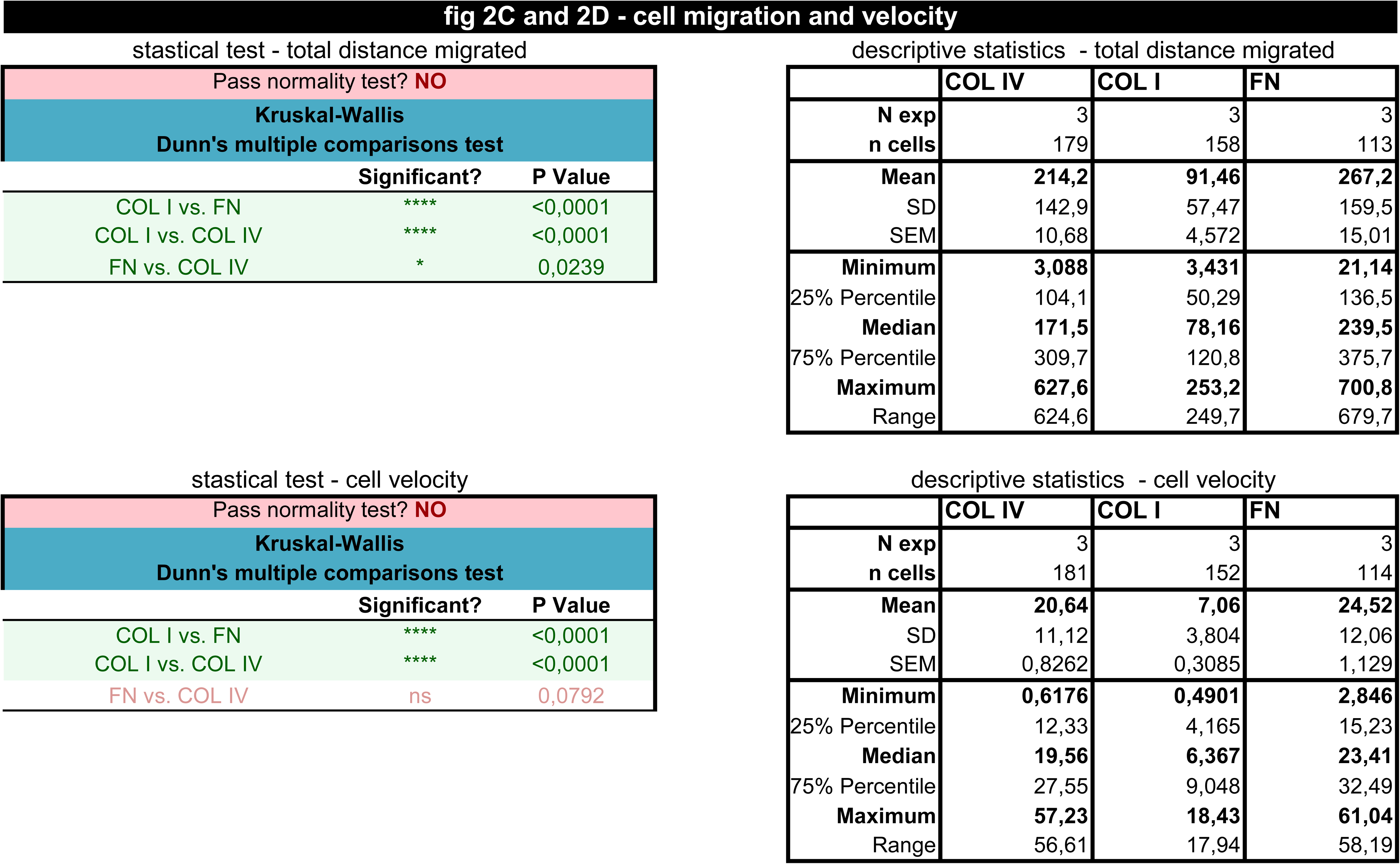
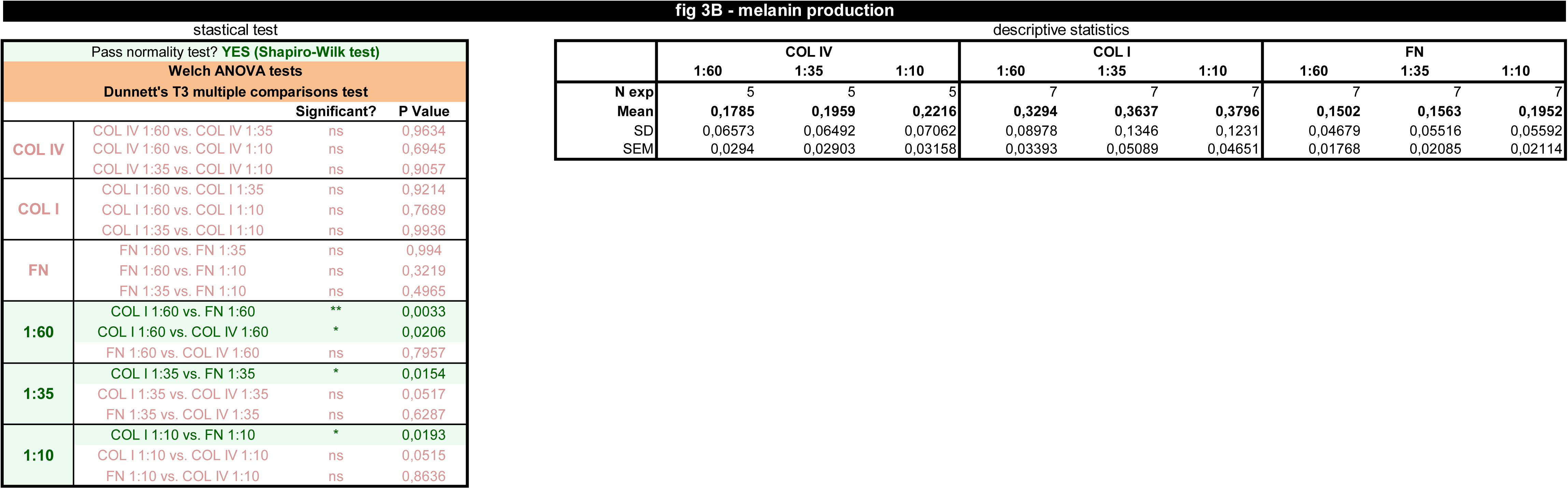
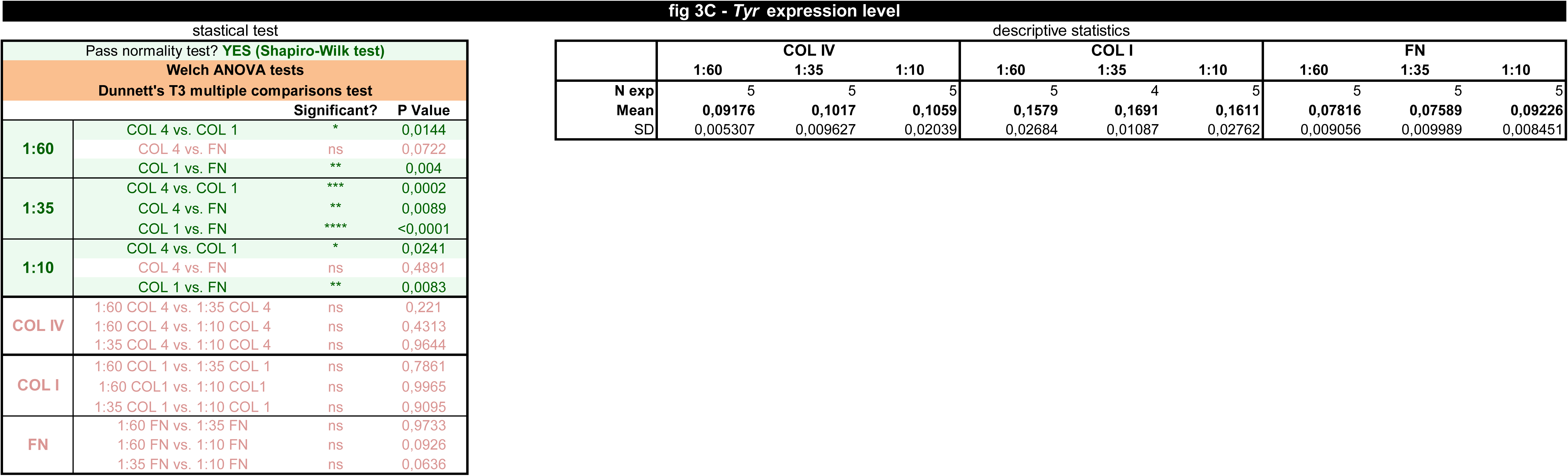
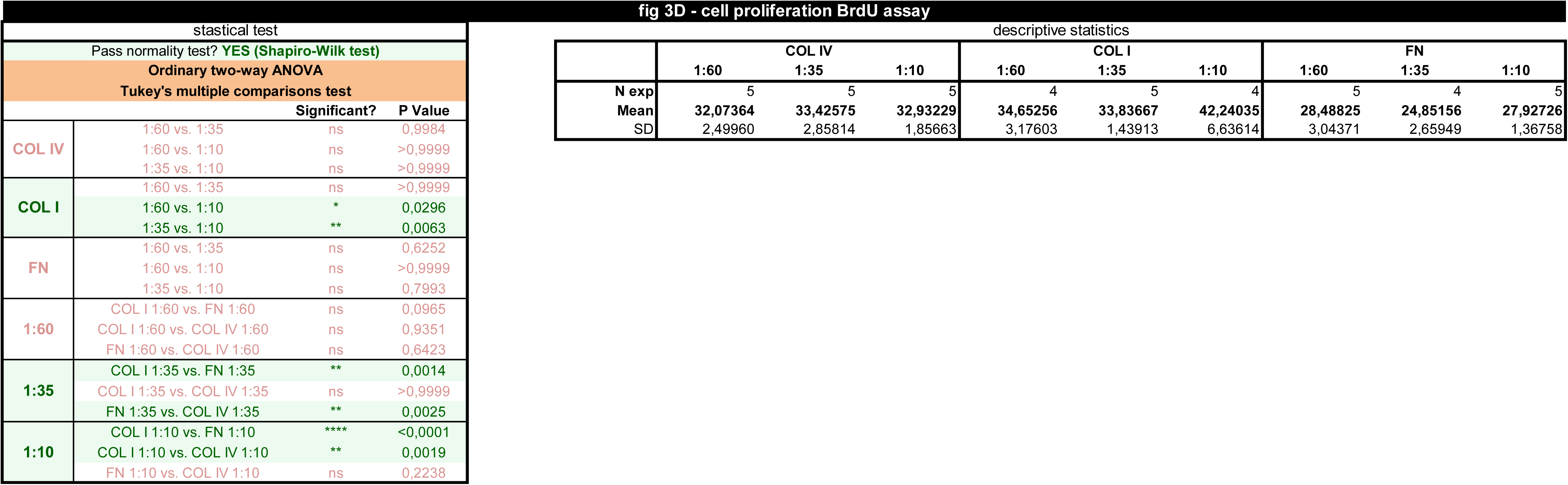
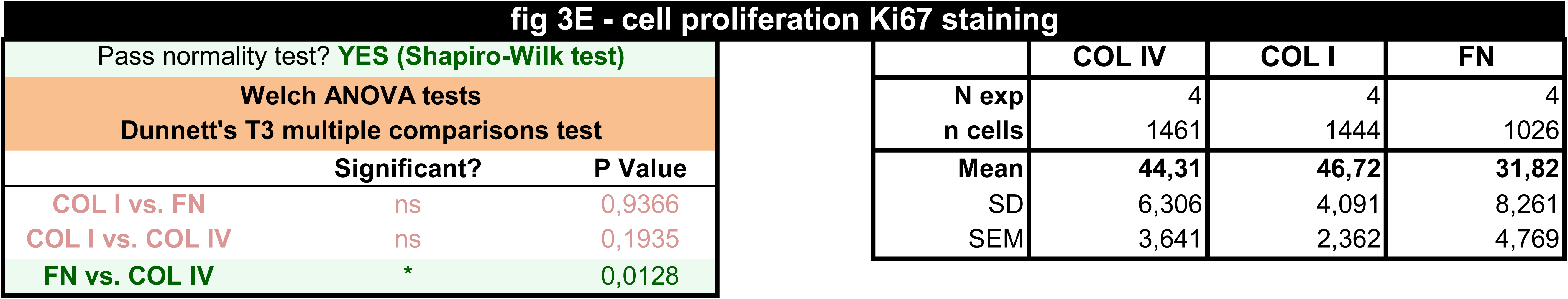
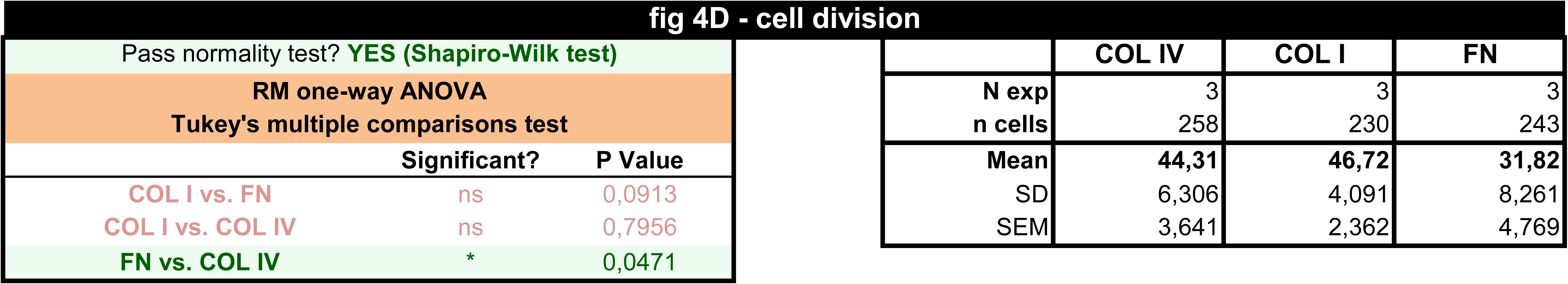
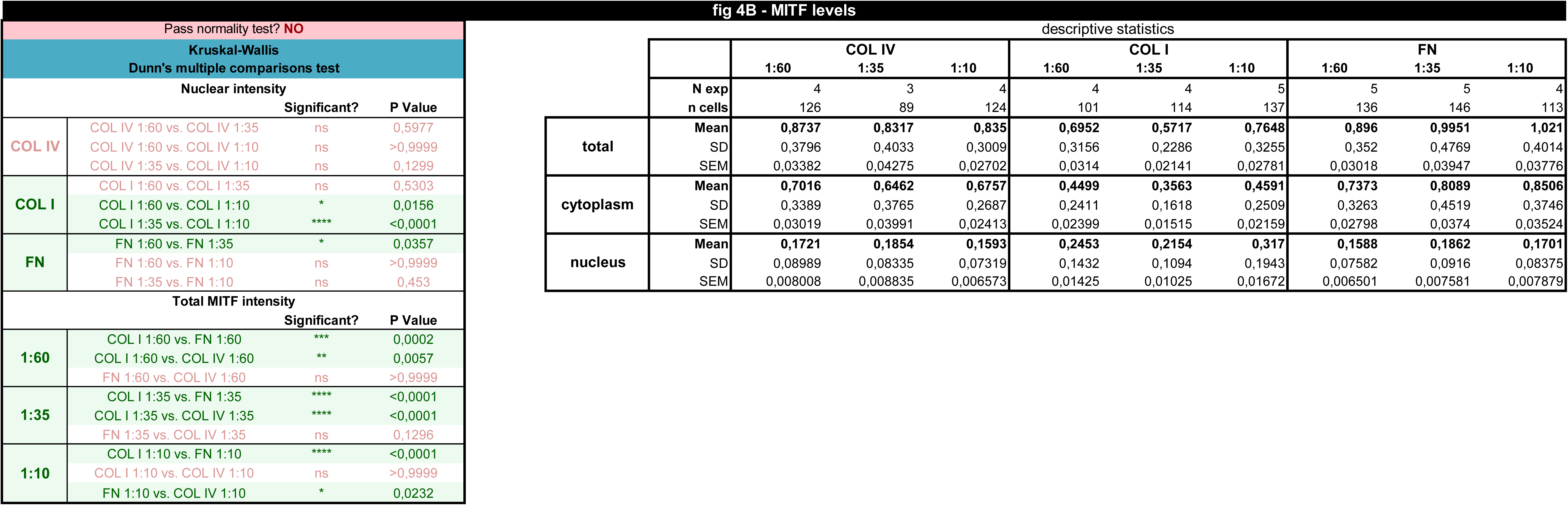
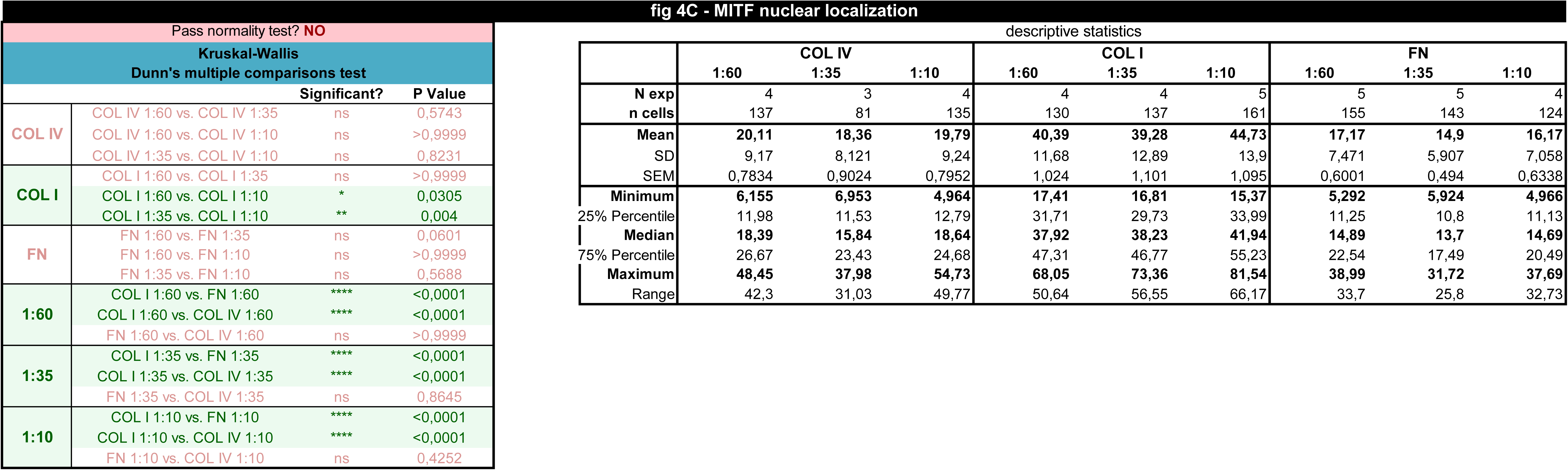
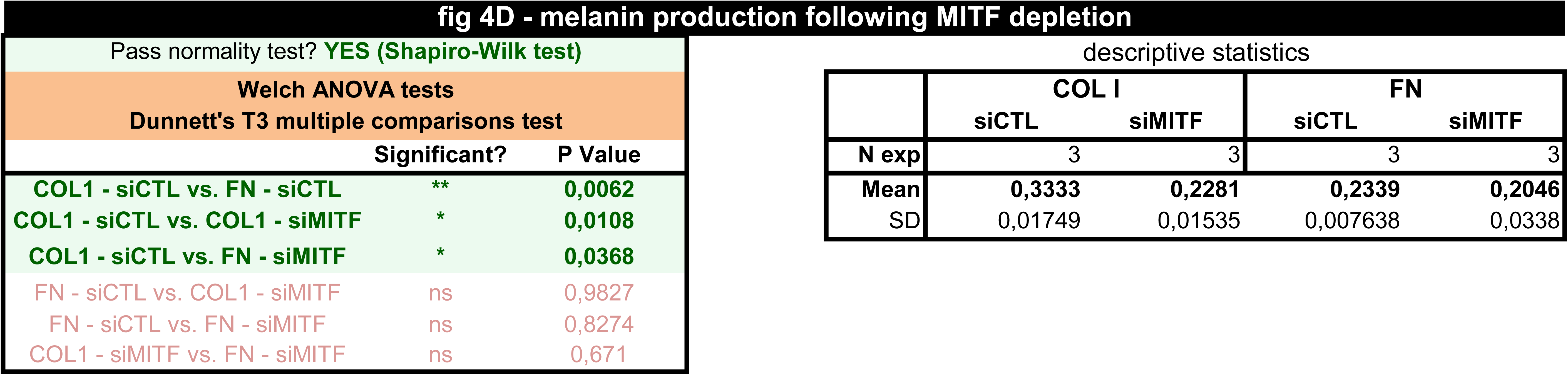
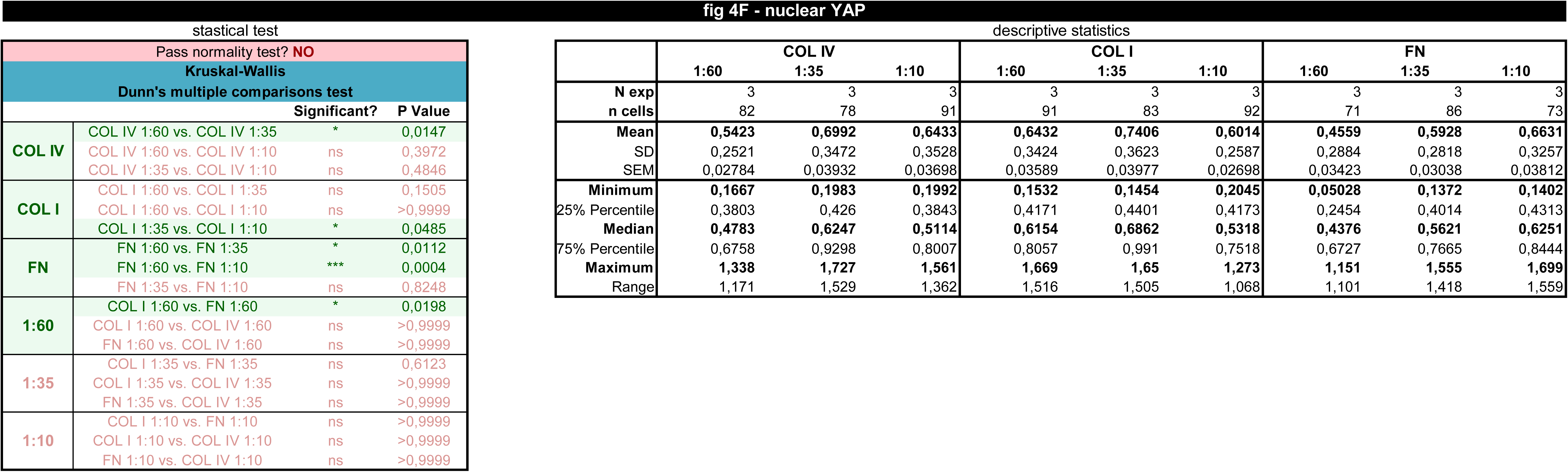
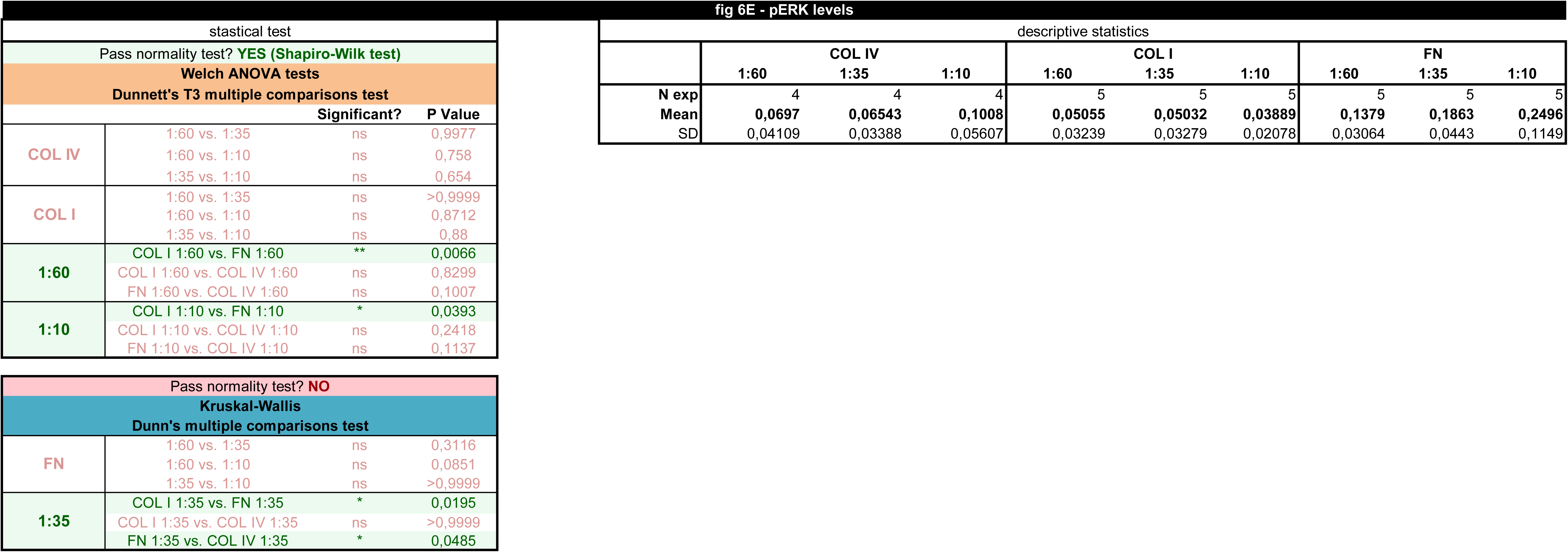
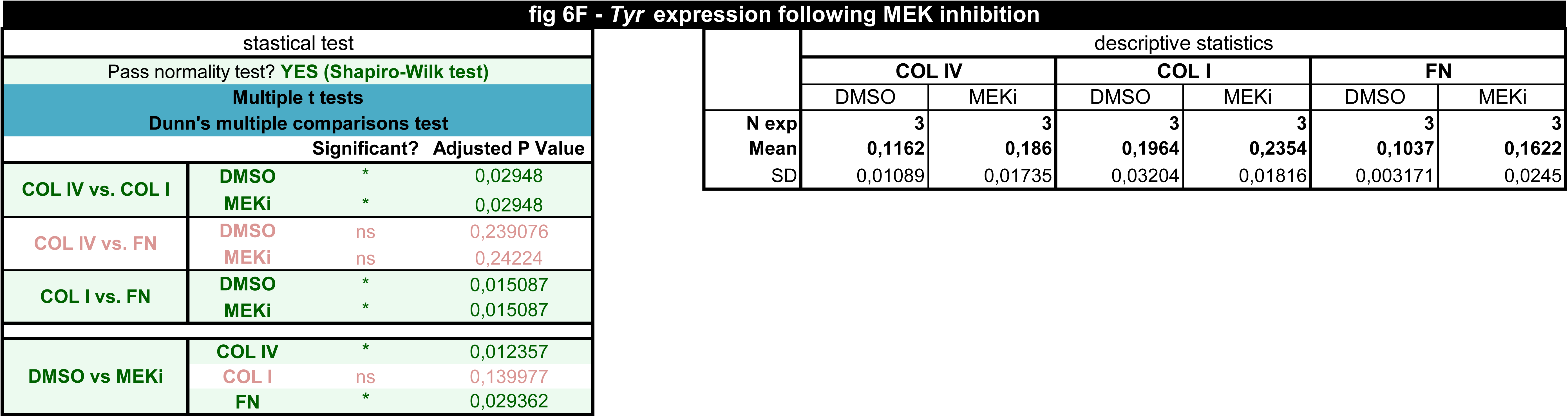
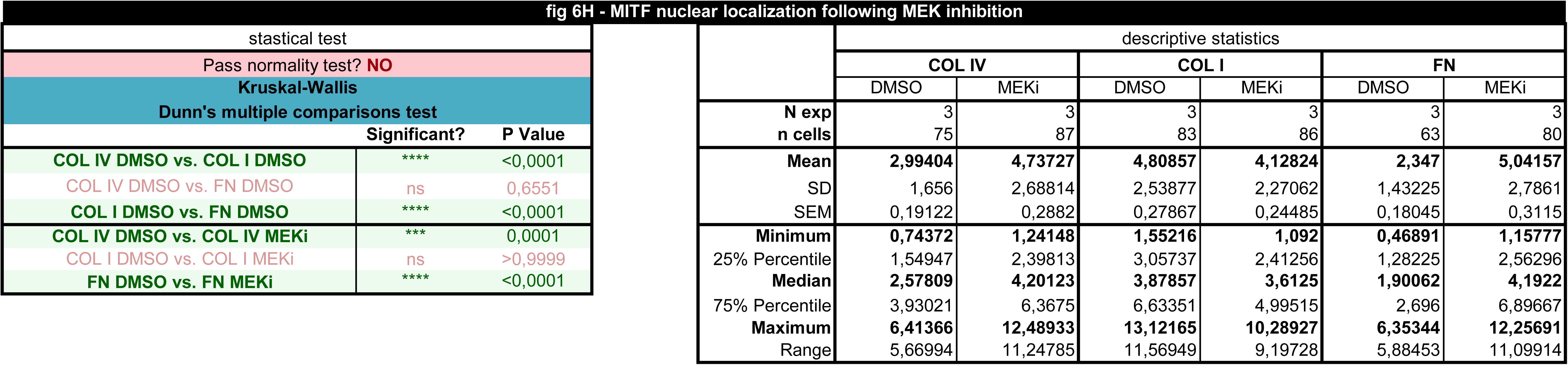
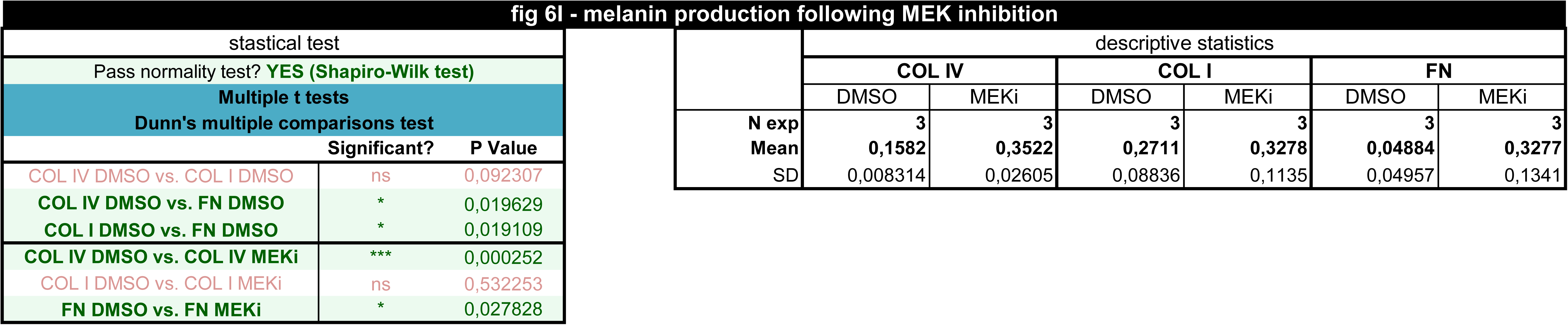
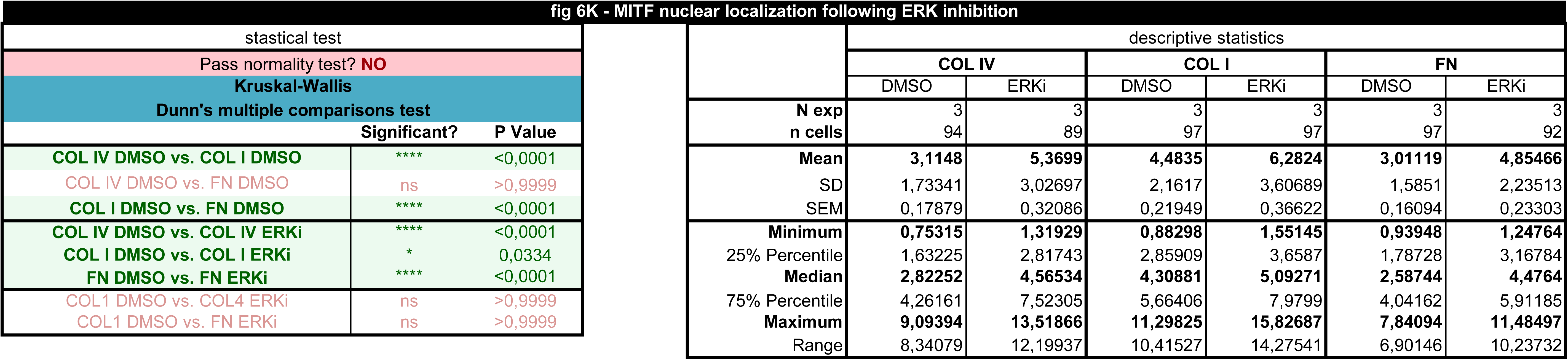
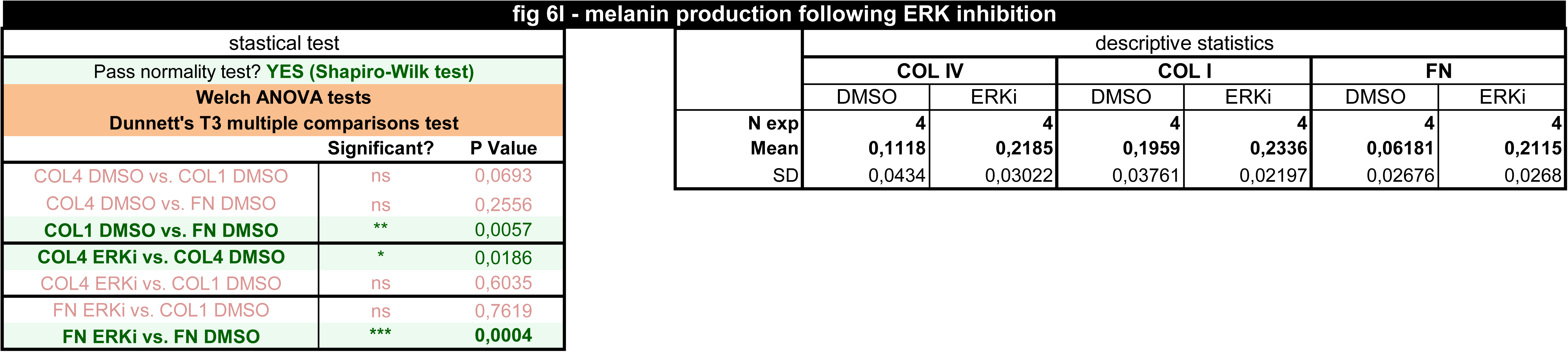
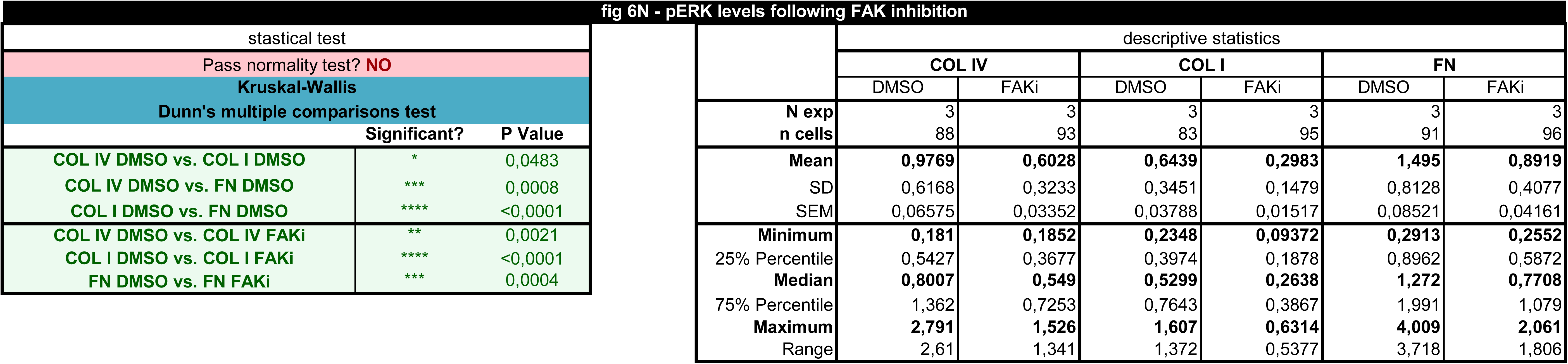
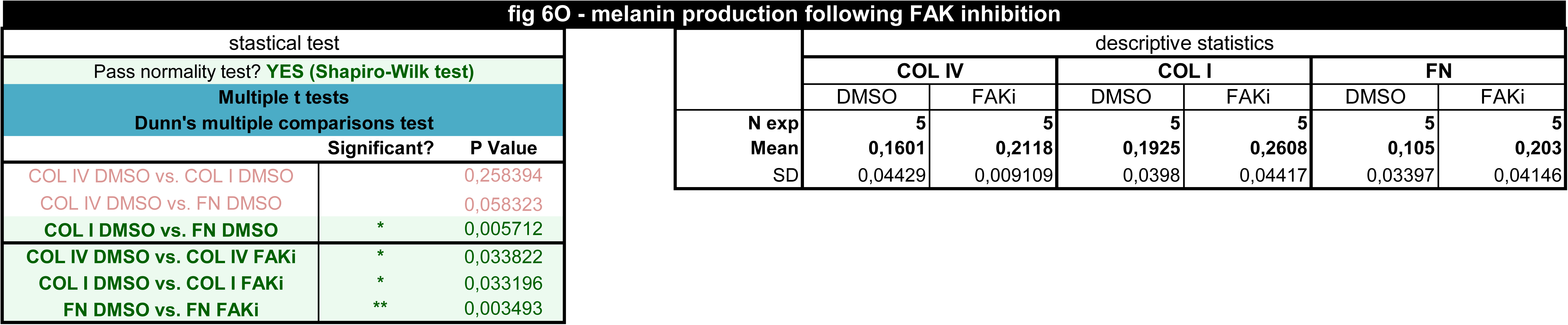
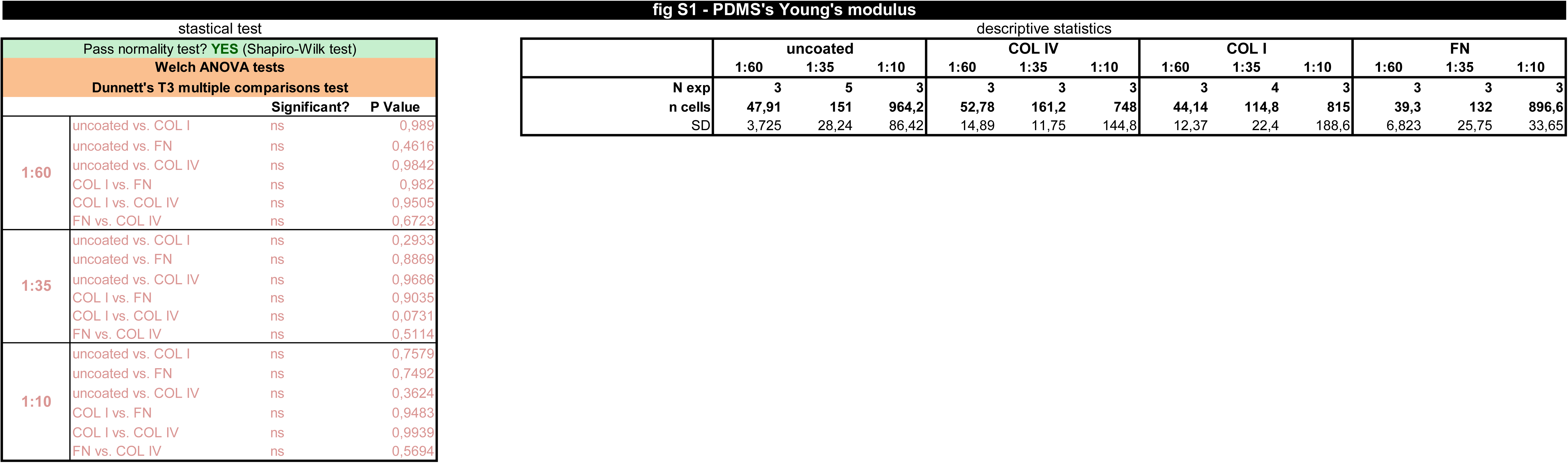
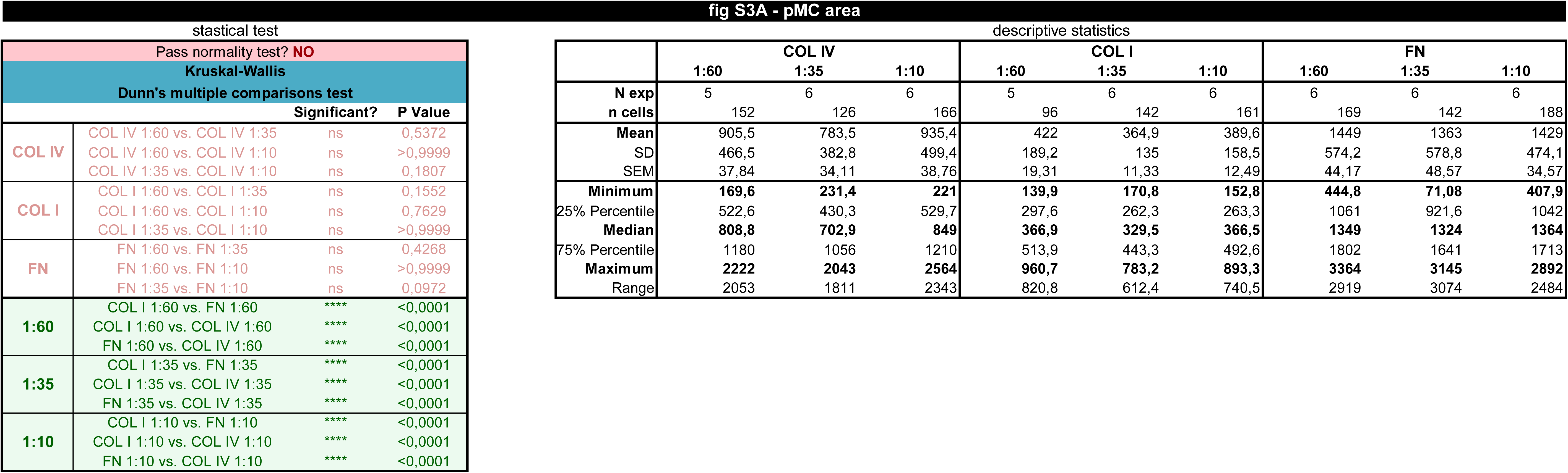
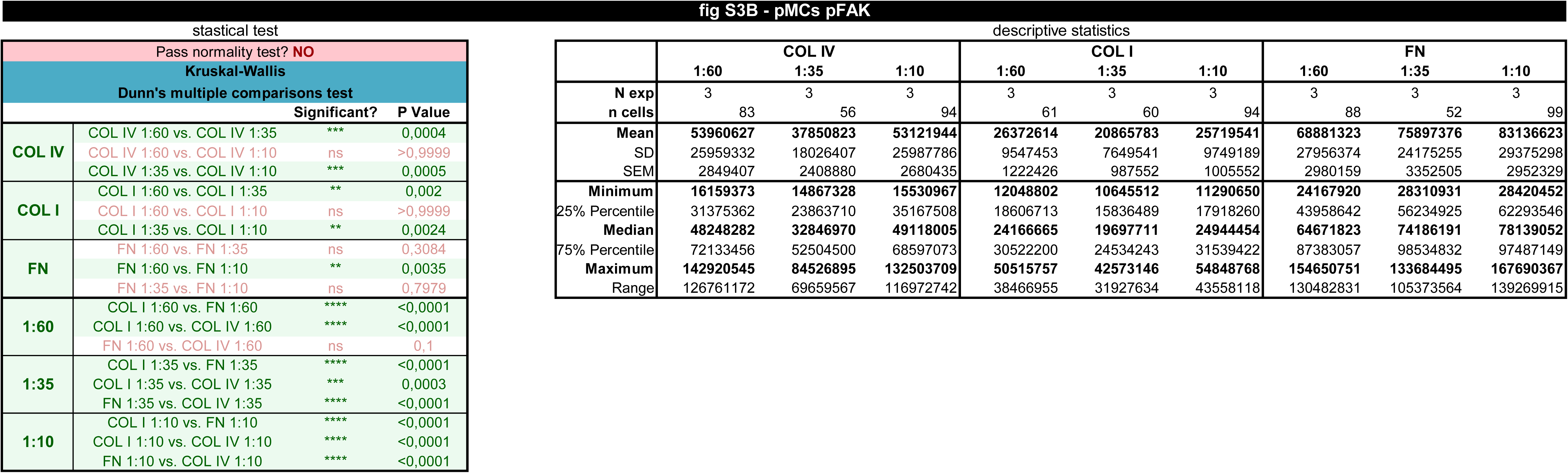
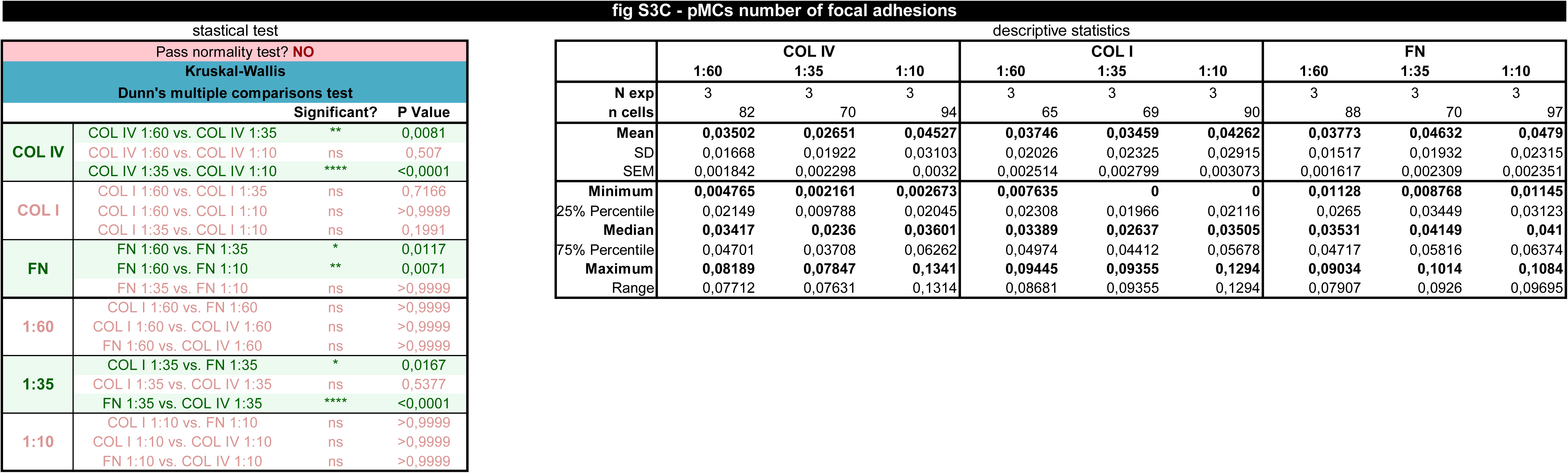
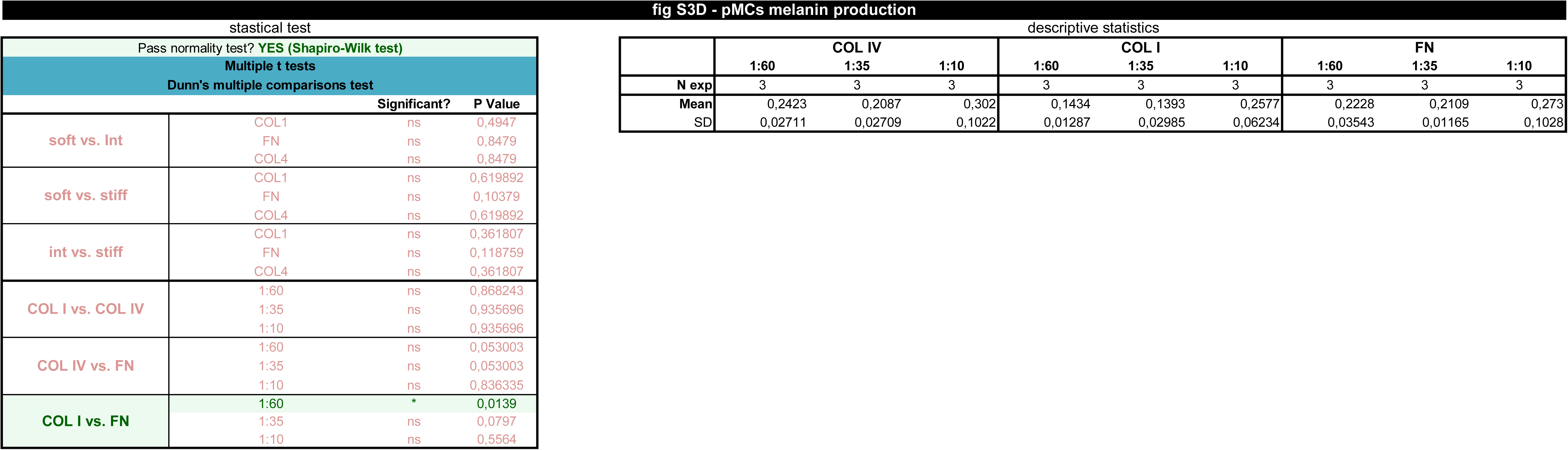
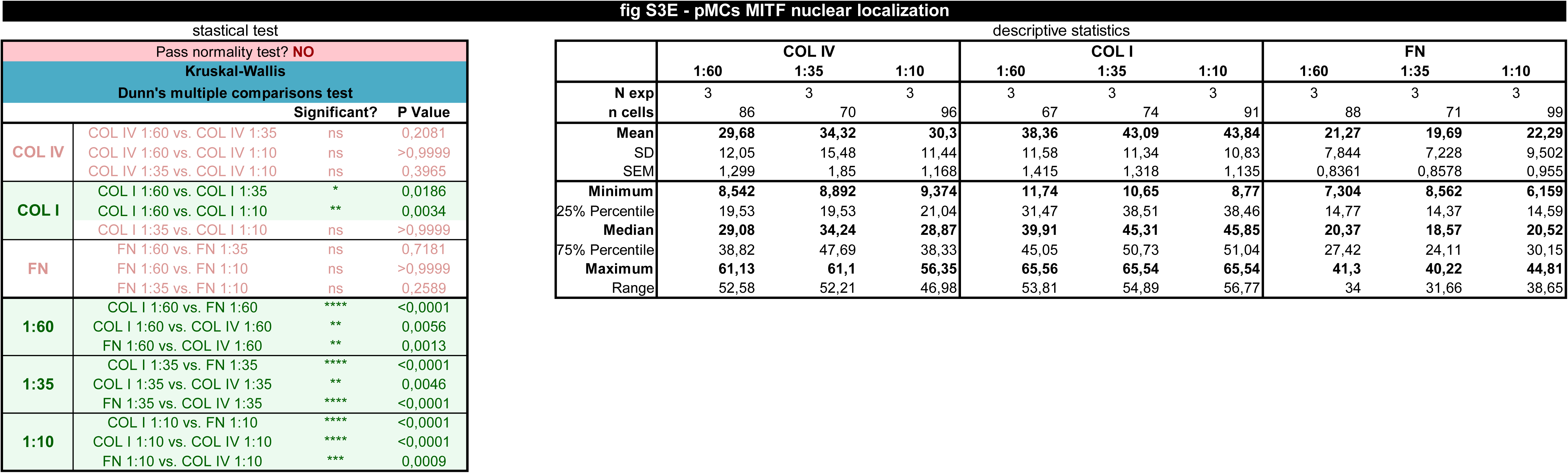
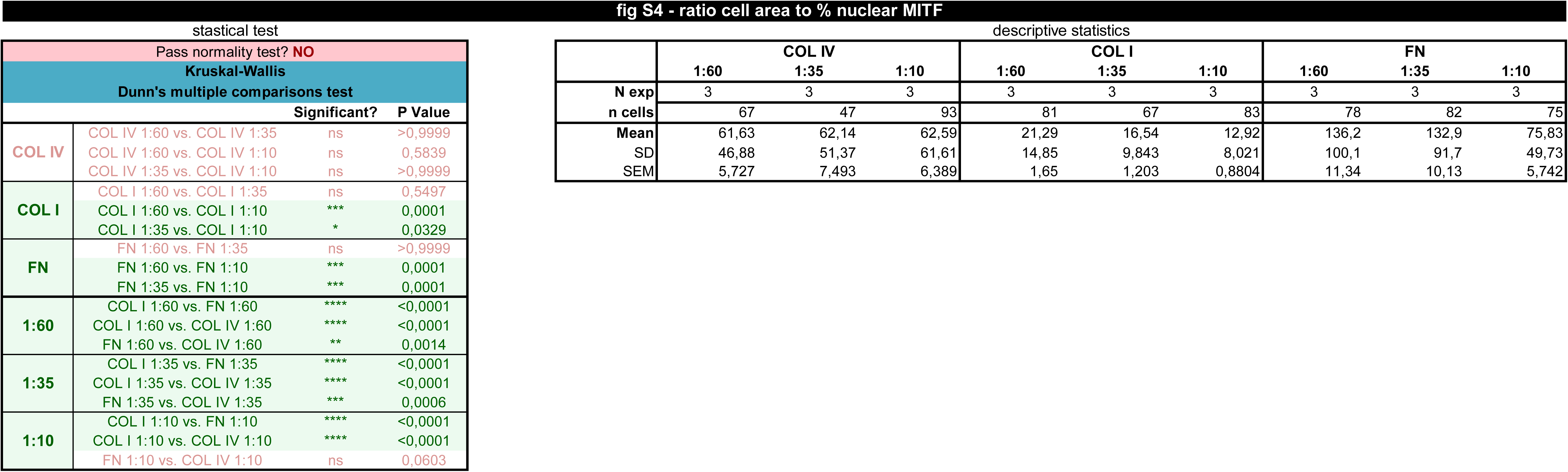
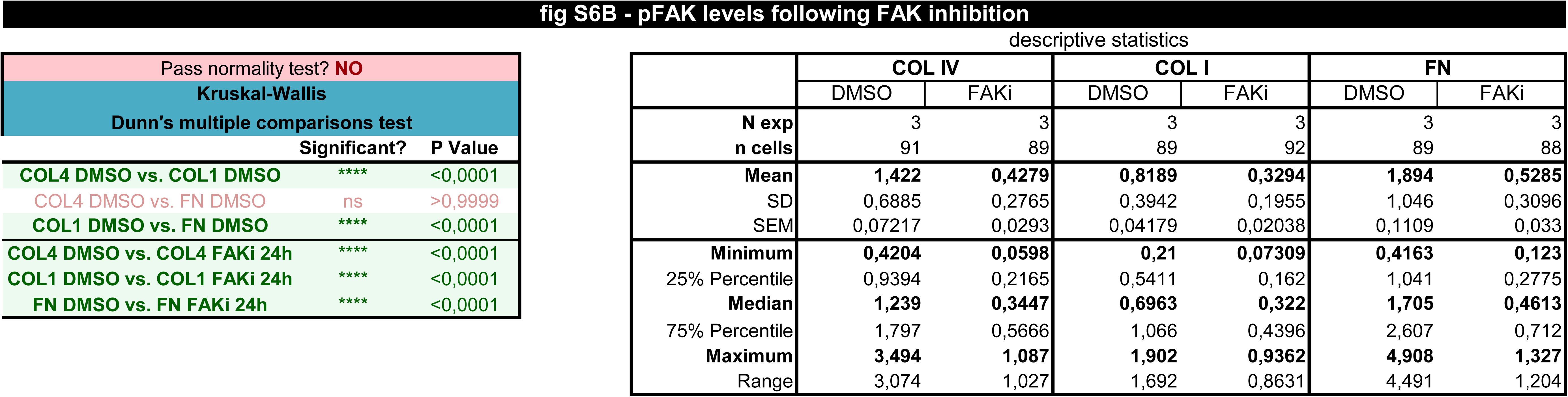
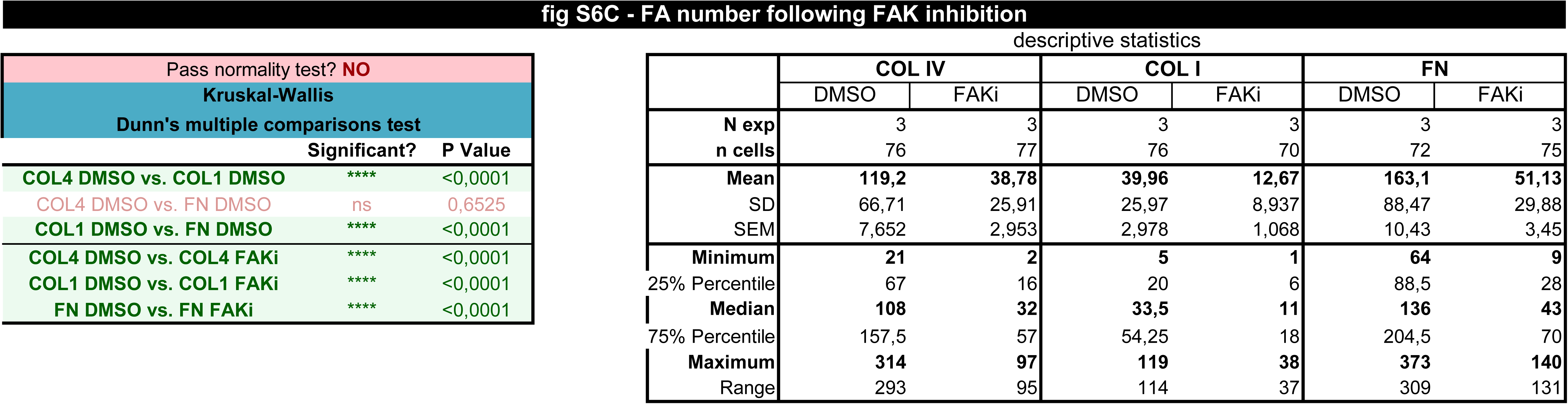

